# High-throughput identification of protein turnover modulators in human neurons

**DOI:** 10.64898/2026.05.06.723176

**Authors:** Joanna Dembska, Anne-Laure Mahul-Mellier, Armelle Tollenaere, Yllza Jasiqi, David M. Suter

## Abstract

Disturbances in protein homeostasis are a defining feature of aging and neurodegenerative diseases. However, current proteomics approaches do not enable screening-scale pharmacological interrogation of protein turnover in post-mitotic human neurons. Here we establish a Mammalian Cell-optimized Fluorescent Timer (MCFT)-based live-cell imaging platform to quantify global protein synthesis and degradation rates in human embryonic stem cell-derived neurons at screening scale. Among 5,897 tested compounds, 199 increased protein synthesis and degradation rates, and three selected compounds upregulated translation-associated genes in human neurons. We then tested their ability to suppress the accumulation of pathogenic α-synuclein aggregates in models of Lewy body–like pathology. All three compounds induced efficient clearance of α-synuclein aggregates in mouse primary neurons, and one compound demonstrated similar efficacy in human dopaminergic neurons. This platform provides a broadly applicable strategy for identifying biomedically-relevant compounds that enhance protein turnover across pluripotent stem cell-derived cellular models.

## Introduction

Proteome homeostasis hinges on the coordination of protein synthesis and protein clearance. The interplay between these processes ensures maintaining a sufficient rate of protein turnover, which preserves the functional proteome required for cellular identity and physiology. This requirement is especially stringent in the brain, where protein degradation is slower than in other tissues, varies across brain cell types, and harbors subcellular organization of translation and protein degradation (*1–6*). Because neurons are post-mitotic and long-lived, they cannot dilute damaged or misfolded proteins through cell division and must therefore preserve proteome integrity over their lifespan, making turnover regulation particularly relevant to neuronal aging and neurodegenerative disease.

Aging progressively destabilizes this balance: protein synthesis is altered by ribosome stalling, translational dysregulation, chronic activation of the integrated stress response, and broader signaling changes linked to senescence, inflammation, and neurodegeneration (*7–11*). In parallel, protein clearance declines in aging with reduced proteasome activity, impaired proteasome stoichiometry, slowed turnover of proteasome subunits, diminished autophagic flux, and reduced protein recycling in the brain and in peripheral organs (*12–15*). Consistent with these changes, protein lifetimes broadly increase across the proteome during aging (*13*, *16*). Beyond aging, protein turnover changes have been implicated in cancer, cardiac hypertrophy, and muscle atrophy, suggesting that turnover failure is a general feature of stressed or diseased cells (*15*, *17–20*). Sequestration of proteostasis factors involved in degradation, trafficking, and folding into protein aggregates can further amplify cellular stress (*21*, *22*), yet direct measurements of protein turnover in neurodegeneration remain limited, leaving it unclear whether the stage-and model-dependent turnover defects extend across neurodegenerative diseases (*14*, *23*, *24*).

These limitations are particularly relevant for Parkinson’s disease (PD) and related synucleinopathies, where defects in protein clearance are caused by defects in both the ubiquitin-proteasome system and lysosomal pathways (*22*, *25–27*). α-Synuclein (aSyn) is cleared primarily by the proteasome under basal conditions. However, pathogenic forms of aSyn interfere with both proteasomal and autophagy pathways (*22*, *26*, *28–30*). Alterations in protein synthesis have also been implicated in PD, with evidence for downregulation in ribosomal and translation-associated pathways in the substantia nigra, as well as altered mRNA stability, ribosomal remodeling, and context-dependent bidirectional changes in protein synthesis (*10*, *31–40*). Because protein synthesis and clearance are tightly coordinated through stress-response and nutrient-sensing pathways (*41–43*), perturbing either process is expected to reshape proteome turnover. This makes turnover both a sensitive systems-level readout of cell state and a candidate therapeutic node: if turnover slows down under proteotoxic stress, then restoring or enhancing it may help re-establish proteome homeostasis. Consistent with this idea, boosting proteasome function, modulating autophagy, restoring chaperone capacity, and tuning translation have all been proposed as protective strategies in neurodegenerative disorders (*27*, *44–49*).

To date, quantitative measurements of protein turnover have largely relied on metabolic labeling and proteomics-based workflows such as dynamic stable isotope labeling with amino acids in cell culture (dSILAC), proteome birthdating or deuterated water (D_2_O) (*1*, *50–55*). However, these approaches typically require substantial amounts of starting material and multiple time points to robustly estimate protein decay rates (*56*). dSILAC is particularly challenging to implement in stem cell-derived human neurons due to demanding culture constraints such as susceptibility to detachment, dependence on specialized media and limited scalability, particularly over prolonged labeling periods (*23*, *57*, *58*). As a result, such approaches have limited throughput and are not suitable for screening-based interrogation of protein turnover modulators in human neurons.

Our laboratory established a Mammalian Cell-optimized Fluorescent Timer (MCFT), which allows to infer protein synthesis and degradation rates of single live cells by fluorescence microscopy (*59*). By benchmarking MCFT-inferred rates to those obtained by SNAP-tag pulse chase labelling, HPG labelling, proteasome activity assays, and dynamic SILAC, we have demonstrated that the MCFT allows robust quantification of perturbations in global protein synthesis and degradation rates (*42*). Here, we leverage the MCFT to establish a quantitative, imaging-based platform to measure protein turnover in individual living human neurons derived from pluripotent stem cells. We combine this system with a high-throughput small-molecule screen to identify modulators of neuronal protein turnover, characterize their impact on neuronal transcriptome and proteome, and demonstrate their ability to modulate pathological features in cellular models of Parkinson’s disease.

## Results

### Human-induced neurons exhibit slow protein turnover

To address the limitations inherent to proteomics-based approaches for measuring protein turnover in neurons, we employed a single-cell, quantitative protein turnover reporter we developed in human embryonic stem cell (hESC)-derived neurons (*42*, *59*). We used a recently engineered hESC line ubiquitously expressing the MCFT (*42*, *59*), hereafter called FT-hESCs, which was inserted into the Citramalyl-CoA Lyase (CLYBL) safe harbor locus (*60*), allowing stable transgene expression in neurons. The MCFT combines two fluorescent proteins with different maturation rates: a fast-maturing green fluorescent protein (super-folder green fluorescent protein (sfGFP)) and a slower-maturing red fluorescent protein (mOrange2). The ratio between the green and red fluorescence signals (G/R ratio) can be used to infer protein synthesis and decay rates (*42*). The MCFT is fused to a PEST sequence to target it directly for proteasomal degradation, thereby providing a measurement of the global cellular protein degradation capacity by the proteasome (*42*, *61*, *62*), and to a SNAP-tag that enables orthogonal measurements of decay rates through pulse-chase labeling (*42*, *63*). To rapidly differentiate hESCs into human neurons, we implemented a doxycycline-inducible, neurogenin-based approach that allows to obtain a nearly pure population of neurons (hereafter, FT-iNGNs) from FT-hESCs by overexpressing neurogenin 1 and 2 (NGN1, NGN2) (*42*, *64*, *65*). Briefly, our FT-hESCs line was engineered using the PiggyBac system to stably introduce doxycycline-inducible NGN1/NGN2 and a rtTA3G expression cassette, generating FT-NGN-hESCs (*42*, *64*, *65*). We then differentiated FT-NGN-hESCs into FT-iNGNs (Fig. 1A) by treating them with dox for 4 days and performed immunofluorescence labeling to validate neuronal differentiation. The vast majority of FT-iNGNs cells were positive for the neuron-specific β3-Tubulin marker (TUBB) and for the dendritic marker Microtubule Associated Protein 2 (MAP2), and negative for the proliferation marker Ki67, confirming the successful differentiation to post-mitotic neurons (Fig. S1A-G). Next, we differentiated FT-hESCs without NGN1/2 overexpression, using well-established neuronal differentiation protocols (*66*) (see Methods section), and we also performed immunofluorescence validation of neuronal markers (Fig. S1G). We then quantified the G/R ratio from the MCFT in undifferentiated FT-hESCs, FT-NGN-hESCs differentiated to FT-iNGNs, or FT-hESCs differentiated to neuron-enriched culture (Fig. 1B-C). As expected, FT-iNGNs exhibited longer MCFT half-lives compared to FT-hESCs (Fig. 1B), similar to those observed in neuron-enriched cultures (Fig. 1B). This indicates that the protein turnover regime of FT-iNGNs resembles long-term differentiated neurons, consistent with previous reports of slower protein turnover in the brain and in neurons compared to other cell types (*1*, *5*, *6*).

**Fig. 1.**
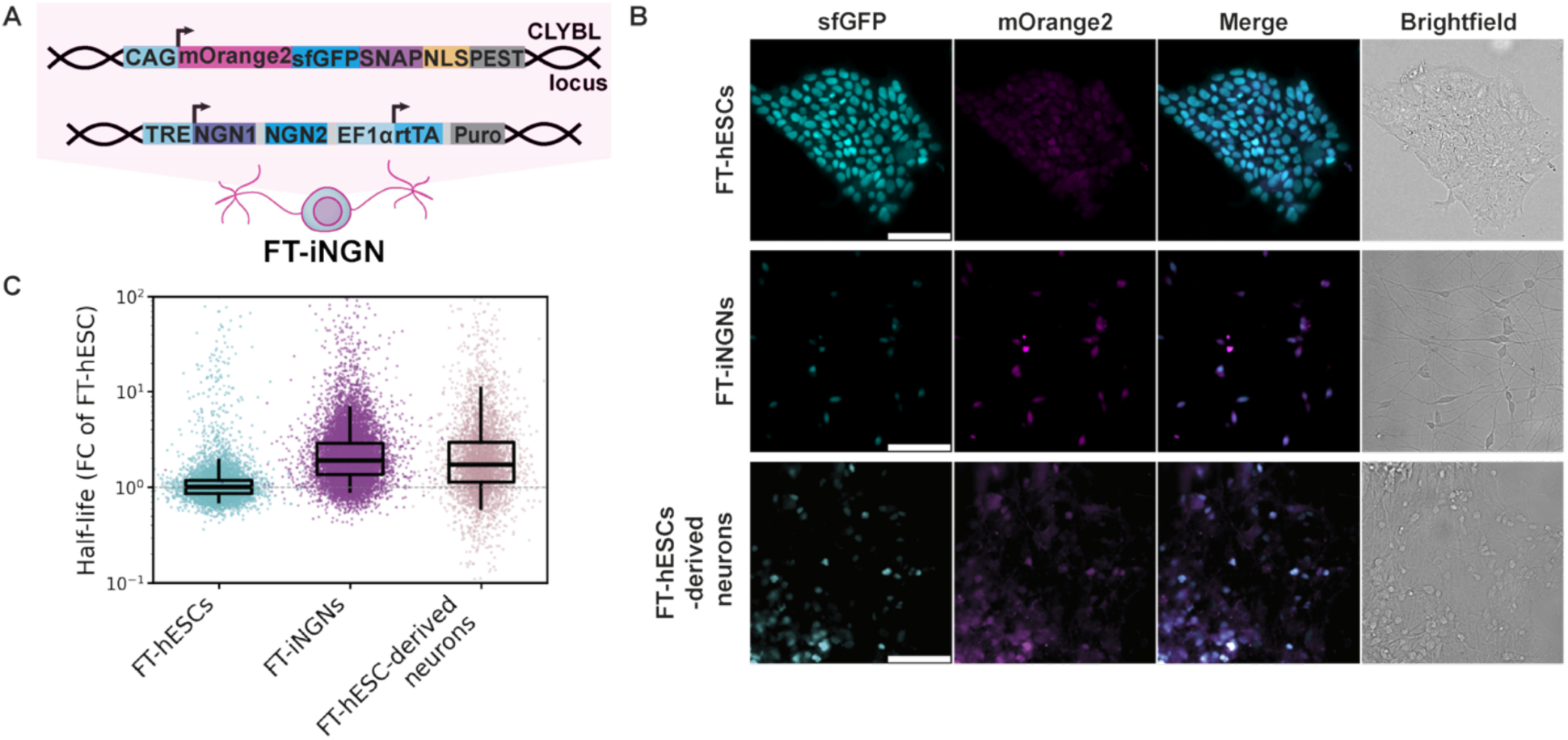
Characterization of fluorescent timer (FT)-expressing undifferentiated FT-hESCs and FT-hESC-derived neurons. **A.** Scheme of FT-iNGNs expressing the Mammalian Cell-optimized Fluorescent Timer (MCFT) and Neurogenin 1 and 2 (NGN1/2). **B.** Representative snapshots of sfGFP (cyan) and mOrange (magenta) channels, merge of both channels, and brightfield of the FT-expressing human cell lines: human embryonic stem cells (FT-hESCs), inducible neurons (FT-iNGNs), neuron-enriched cultures derived from FT-hESCs, respectively. Scale bar: 100 µM. **C.** Half-lives of the MCFT in FT-hESCs (cyan), FT-iNGNs (magenta), FT-hESC-derived neuron-enriched culture (pink), computed from the green to red (G/R) fluorescence ratio (see Methods), normalized to the median of FT-hESCs. Each point represents a single cell (FT-hESCs: N= 7390; FT-iNGNs: N=11286, FT-hESC-derived neuron-enriched culture: N= 4429). Boxes: interquartile range; horizontal line: median; dashed line: median of FT-hESCs; vertical lines: 5^th^-95^th^ percentiles.

### High-throughput screening identifies protein turnover modulators in human neurons

We next set out to identify small molecules that enhance protein turnover in post-mitotic FT-iNGNs. We performed an imaging-based screen to obtain quantitative single-cell measurements of green and red fluorescence (Fig. 2A). We pre-dispensed compounds into a 384-well plate with an acoustic liquid handler, added FT-iNGNs pre-mixed with laminin using an automatic cell dispenser, and we imaged the plates with a high-throughput fluorescence microscope after 24 h (see Methods). We then determined a screening window coefficient (z’-factor) to assess the quality and variability of the data (*67*). Since positive controls for increased protein turnover validated for our specific experimental system were not readily available, we first tested control conditions versus cycloheximide (CHX) treatment, which lowers protein turnover rates (*42*). We compared 194 wells of FT-iNGNs-treated with vehicle (DMSO) to 194 wells treated with 20 µM CHX (Methods). The screening window coefficient was determined as z’=0.68 (Fig. 2B), indicating that the separation window of our assay is of high quality (*67*). We then designed the screening itself so that each plate included vehicle-treated control wells and CHX-treated wells, allowing calculating the z’-factor to assess plate quality and perform per-plate and per-batch normalization. Only plates with a z’-factor above 0.25 were used for further analysis (see Table S1).

**Fig. 2.**
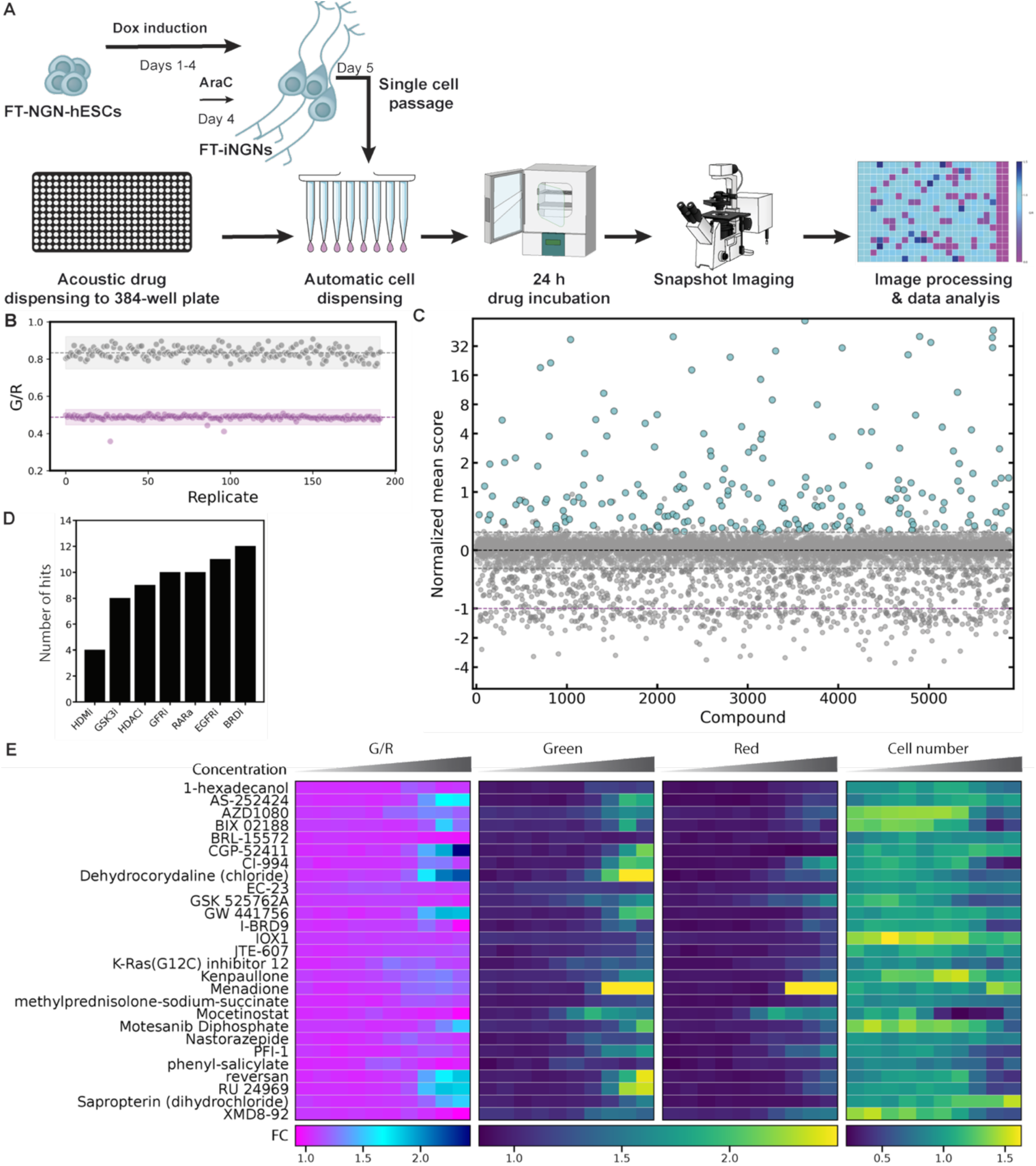
Small-molecule screening reveals protein turnover modulators in human neurons. **A.** Scheme of the screening workflow. **B.** Screening window coefficient (z’-factor): FT-iNGNs were cultured following the screening workflow from (**A**), treated for 24 h with a vehicle (DMSO, gray) and a control compound (20 µM CHX, magenta). Each dot represents a replicate (N=194 wells, at least 800 cells per replicate were measured), dashed line: mean value, colored window: 3 standard deviation values in each direction. **C.** Normalized inverted mean values of drugs tested in the primary screen (N=2). Compounds (gray) were annotated as hits (cyan) when they scored above 3 standard deviations of the DMSO control. The inverted values are displayed, where CHX =-1 (dashed magenta line) and DMSO = 0 (dashed black line). At least 800 cells per compound per replicate were measured. **D.** Barplot of the number of identified hits in the most prevalent classes in the primary screen. HMDi: Histone methyltransferase inhibitors; GSK3i: Glycogen Synthase Kinase 3 inhibitors; HDACi: Histone Deacetylase inhibitors; GFTi: Growth factor tyrosine kinase inhibitors (targeting either PDGFR and/or VEGFR and/or FGFR); RARa: Retinoic Acid Receptor agonists; EGFRi: Epidermal growth factor receptor inhibitors; BRDi: Bromodomain inhibitors. **E.** Heatmap of the fold change (FC) of the mean G/R ratio, mean green, mean red fluorescence, cell number of the compounds tested in the secondary screen (excluding compounds removed due to precipitation or imaging artifacts), normalized to the vehicle (DMSO) (N=2; at least 800 cells per compound per replicate were measured).

In total, we screened 5897 compounds coming from three libraries: an in-house kinase inhibitors library (Supplementary file S1), a drug repurposing library (Supplementary file S2), and the Prestwick Chemical Library® (Prestwick Chemicals). These libraries contained mostly FDA-approved drugs and compounds at different stages of pre-clinical and clinical studies, the majority of which have known mechanisms of action (MOA). Each compound was tested in duplicate, using a 10 µM concentration and a 24 h incubation time. Hits were defined as compounds that increase the G/R ratio by at least three times the standard deviation of the mean in DMSO-treated cells for both replicates. The linear correlation coefficient R^2^ of duplicates was 0.722 (Fig. S2A), excluding compounds from one plate that did not pass the quality assessment of the z’-factor. As expected, known proteasome and protein synthesis inhibitors were among the 900 compounds that decreased the G/R ratio (Fig. S2B), and we found 199 compounds that increased the G/R ratio (Fig. 2C).

Hits from the primary screen were manually filtered to remove false positives, such as compounds that precipitated or caused imaging artifacts. We also removed compounds that caused more than a 2.5-fold decrease in cell numbers based on the number of segmented cells, or those that had similar chemical structures, retaining only the most active compounds. We then grouped the remaining hits by MOA. The most frequent MOA classes included growth factor receptor kinase inhibitors (PDGFR/FGFR/VEGFR and EGFR inhibitors), histone deacetylase inhibitors (HDACi), bromodomain inhibitors (BRDi), glycogen synthase kinase 3 inhibitors (GSK3i), retinoic acid receptor agonists (RARa), and histone demethylase inhibitors (HDMi), represented by 8 to 12 compounds in the most frequent classes and 4 compounds in the HDMi class (Fig. 2D). For the most frequent MOA classes, we selected three compounds based on activity, whereas for RARa and HDMi we retained only the single most active compound because of their overall lower activity towards enhancing protein turnover. The final set was complemented with the most active compounds from less frequent MOA classes. Using these criteria, we selected 47 compounds for the secondary screen and performed dose-response curves at 10 concentrations ranging from 3 nM to 40 µM.

The experimental and analytical workflows were kept as in the primary screen. All plates had a z’-factor above 0.5, indicating sufficient quality (Table S1). We again manually reviewed the images to exclude compounds with precipitation or imaging artifacts. For the remaining compounds (Table S2), we compared G/R ratios, changes in green and red fluorescence, and cytotoxicity (Fig. 2E). By fitting our data to a sigmoidal function, we calculated a half-maximal effective concentration (EC50). We only kept compounds that i) increased the G/R ratios by at least 20%, ii) did not reduce cell number, iii) had an EC50 below 30 µM, iv) and showed robust dose-response fitting (R² > 0.55). Using these criteria, together with a literature-based evaluation of known biological properties and limited prior links to protein turnover, proteostasis, and neurodegeneration, we selected three compounds for further investigation: AS-252424 (PI3K inhibitor), CGP-52411 (EGFR inhibitor) and CI-994 (HDAC inhibitor) with EC50 of 10, 23 and 3.3 µM, respectively (Fig. S2C). Interestingly, for all three compounds, the G/R ratio increase was driven mainly by higher green fluorescence (Fig. S2D), while red fluorescence (Fig. S2E) was slightly reduced upon CGP-52411 treatment and increased less strongly than green fluorescence upon AS-252424 and CI-994 treatment, without reduced cell numbers (Fig. S2F), except for CI-994 above 10 µM. Increases in green fluorescence were not due to increased nucleus size (Fig. S2G). The lack of changes in cell number (Fig. S2F), nuclear size (Fig. S2G), or cellular morphology (Fig. S2H) suggests that drug treatment did not induce overt changes in the cellular state relative to the pre-treatment post-mitotic condition (Fig. S1B). This combined increase of green fluorescence and G/R ratio indicates that the compounds increased both protein synthesis and protein degradation. Interestingly, numerous compounds belonging to the same mechanistic classes (i.e., PI3K, EGFR, and HDAC inhibitors, respectively), either reduced the G/R ratio or had no effect (Fig. S2I-K).

### Protein turnover modulators enhance expression of ribosomal protein genes

We next examined the impact of a 24 h treatment with selected compounds on the neuronal transcriptome. We performed RNA sequencing on iNGNs cells treated with 20 µM AS-252424, 20 µM CGP-52411, and 5 µM CI-994 alongside vehicle-treated (DMSO) controls. AS-252424 treatment resulted in minor changes (Fig. 3A), whereas CGP-52411 led to broader changes in the transcriptome (Fig. 3B). In both treatments, most top upregulated genes were involved in translation, such as ribosomal proteins (RPL39, RPS3A), translation initiation factors (EIF3E, EIF3L) and various Small nucleolar RNA (snoRNA) (SNORA63, SNORD97, SNORA12, SNORA49, SNORD116-18, SNORD15B), previously reported to play a role in rRNA/tRNA modification, pre-rRNA processing, ribosome biogenesis, and translation control (*68–71*). In contrast, CI-994 triggered a large transcriptional response, with 6004 downregulated and 1880 upregulated genes (Fig. 3C), consistent with the known activity of CI-994 as a class I histone deacetylase (HDAC) inhibitor (*72*). While HDAC inhibition is generally expected to increase gene expression through enhanced histone acetylation, the large fraction of downregulated genes observed here suggests that after 24h of treatment, secondary regulatory effects likely dominate the transcriptional landscape, as shown before in the mouse brain after a single dose of CI-994 treatment (*73*). Next, we asked which biological pathways were most altered by each treatment using gene ontology enrichment analysis of the differentially expressed genes (DEGs). The majority of the top 15 enriched Reactome terms among genes upregulated by AS-252424 were associated with translation-related programs, including translation initiation and ribosome assembly (Fig. 3D), but no pathways were enriched among downregulated DEGs. Similarly, upregulated genes in CGP-52411-treated iNGN cells were mainly enriched for translation initiation, amino acid transport, and ribosome biogenesis (Fig. 3E), while pathways linked to DNA methylation and chromatin remodeling were depleted (Fig. S3A).

**Fig. 3.**
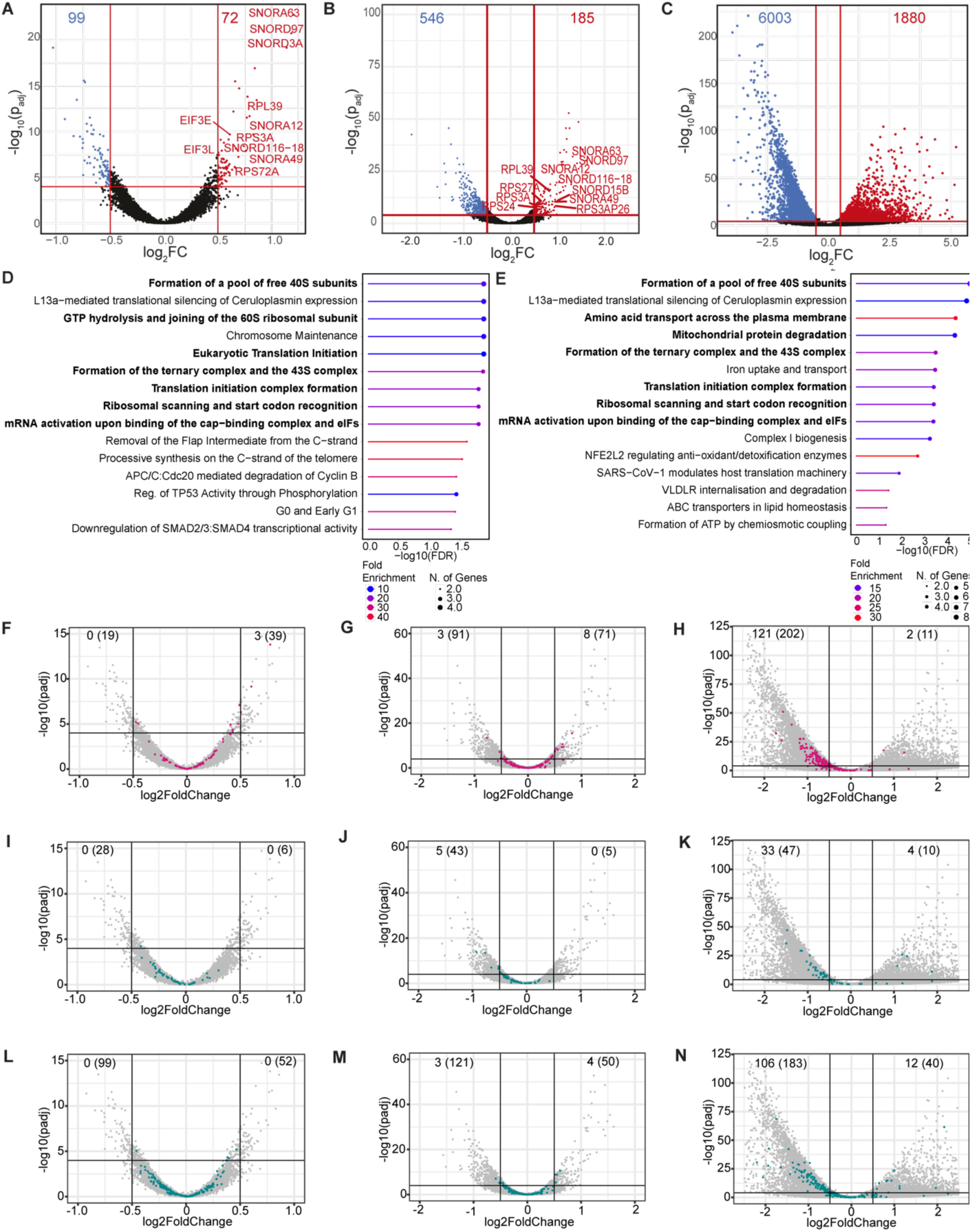
**AS-252424, CGP-52411 and CI-994 remodel the neuronal transcriptome.**Differential gene expression analysis for iNGNs treated with AS-252424 (**A**), CGP-52411 (**B**) and CI-994 (**C**). Each point is the average over biological duplicates of the log2 fold-change (FC) in RNA levels relative to the vehicle treatment (DMSO). **D**-**E**. Reactome Knowledgebase enrichment analyses of the top 15 significantly enriched pathways upregulated in AS-252424-treated (**D**) and in CGP-52411-treated cells (**E**), in bold translation-related terms. **F**-**N**. Differential expression for specific classes of genes (colored dots); (**F-H**): cytoplasmic ribosomal proteins, (**I-K**): proteasomal degradation, (**L-N**) autophagosomal degradation for cells treated with AS-252424 (**F,I,L**), CGP-52411 (**G,J,M**) and CI-994 (**H,K,N**). Number of significantly downregulated or upregulated genes (log2FC>|0.5|, padj<0.0001) as well as genes with log2FC>|0.1| (number between brackets) are displayed on the upper corners of each plots.

In contrast, CI-994 caused broad suppression of pathways involved in DNA methylation, signaling and others, alongside relative enrichment of anabolic processes and long-term potentiation (Fig. S3B–C). We next examined whether pathways related to protein processing and proteostasis were affected. For AS-252424 and CGP-52411, cytosolic ribosomal protein genes, showed an upward trend (Fig. 3F-G), whereas proteasome-associated genes tended to be reduced (Fig. 3I-J). Autophagy-related genes and mTOR pathway components changed in both directions but with small effect sizes (Fig. 3L-M; Fig. S3G–H). In line with these modest transcriptional shifts, pathways closely tied to the known targets of AS-252424 (PI3K) and CGP-52411 (EGFR/PI3K) also showed bidirectional, low-magnitude changes, arguing against strong pathway suppression at this 24 h time point (Fig. S3D-F). In contrast, CI-994 broadly decreased expression across the major turnover-related pathway modules examined, consistent with global transcriptomic remodeling (Fig. 3H,K,N; Fig. S3I). We verified that neurogenesis-related genes were not massively dysregulated upon treatment with AS-252424, CGP-52411 and CI-994, excluding a change in cell identity (Fig. S3J-L), which is in line with the image-based observations from the secondary screen.

We next performed label-free quantitative (LFQ) proteomics on iNGNs using the same experimental workflow as for RNA-seq. Changes in the proteome remained much more limited than in the transcriptome after 24 h of treatment for all three compounds (Fig. S4A–C). Nevertheless, ribosomal proteins displayed a consistent increase across all conditions (Fig. 4A-C). In contrast, components of the protein degradation machinery, including proteasomal (Fig. 4D-F) and autophagy-related proteins (Fig. 4G-I), were largely unchanged or showed a slight downward trend, indicating a minimal response in protein abundance. Proteins related to mTOR signaling and mTOR positive and negative regulators were largely unchanged (Fig. S4D–F). Taken together, our data suggest that the observed modulation of protein synthesis is driven by increased levels of the core translational machinery (Fig. 4J-L), whereas the increase in protein degradation is more likely mediated by changes in the assembly/activity of the proteasome.

**Fig. 4.**
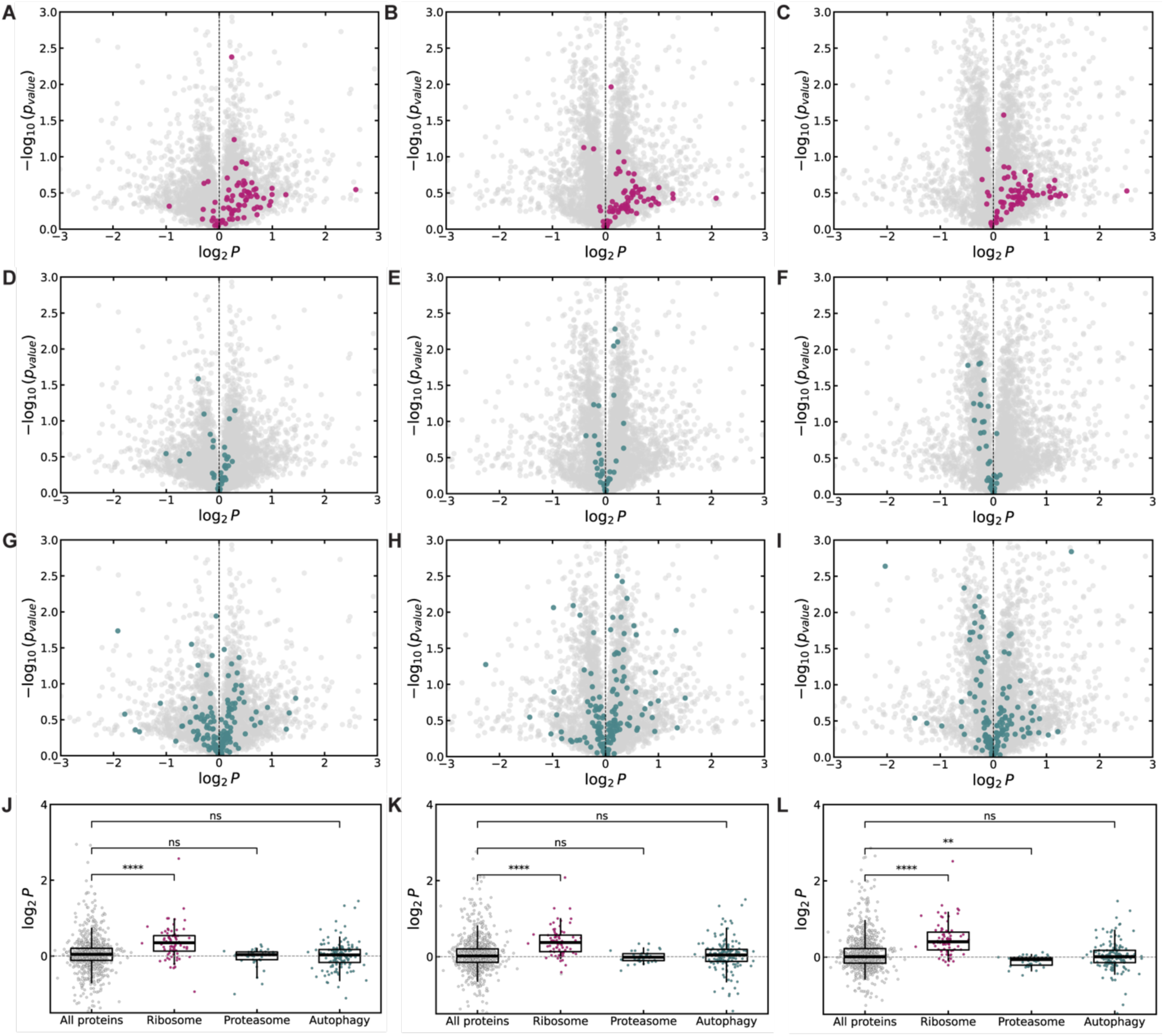
AS-252424, CGP-5241 and CI-994 increase levels of ribosomal proteins. Differential expression for specific classes of proteins (colored dots). (**A-C**): ribosomal proteins, (**D-F**): proteasomal proteins, (**G-I**): and autophagy-related proteins for cells treated with AS-252424 (**A, D, G**), CGP-52411 (**B, E, H**) and CI-994 (**C, F, I**). **J-L**. Comparison of differential expression for specific classes of proteins against all other proteins for AS-252424 (**J**), CGP-52411 (**K**) and CI-994 (**L**). Each dot represents a mean of N=3; P: fold-change of protein level; boxes: interquartile range; horizontal line: median; dashed line: median of all proteins; vertical lines: 5^th^-95^th^ percentiles; ** p<0.01; **** p<0.0001; ns - non-significant.

### Protein turnover modulators suppress seeding initiation and pS129 pathology in a neuronal aSyn aggregation assay

Given that impaired protein turnover contributes to the accumulation of misfolded proteins in neurodegenerative diseases, we next asked whether the identified modulators could limit the formation of pathological protein aggregates. We focused on aSyn, a presynaptic protein that is normally cleared by proteostasis pathways but can misfold and assemble into intracellular aggregates. These aggregates enriched in phosphorylated aSyn at residue S129 (pS129) are a defining pathological hallmark of synucleinopathies, including PD, multiple system atrophy, and dementia with Lewy bodies (LB) (*74*). Their progressive accumulation is closely associated with neuronal dysfunction and degeneration, making aSyn aggregation a relevant system to assess the functional impact of modulating protein turnover. To model seeded aSyn aggregation, aSyn preformed fibrils (PFF) were generated and characterized by Coomassie-stained SDS-PAGE, Thioflavin T fluorescence assay and transmission electron microscopy, confirming that both mouse and human aSyn PFF consist of β-sheet-rich amyloid fibrils and are efficiently fragmented by sonication into short seeds suitable for neuronal seeding assays (Fig. S5A-H).

Having characterized the effects of the compounds on transcription and translation in FT-iNGNs, we next examined their impact on seeded aSyn aggregation. Because this process requires sufficient endogenous expression of aSyn, we first assessed aSyn levels in FT-iNGNs by WB (Fig. S5J). aSyn was not detected in these neurons, and they could therefore not be used for the aggregation assay. We therefore used primary mouse neurons, a well-characterized model widely used in PD research to study seeded aSyn aggregation and screen for modifiers of pS129-aSyn pathology (*30*, *75–79*). Following exposure to mouse aSyn PFFs, these neurons reproducibly develop abundant aSyn pathology within 10 days, including the progressive aggregation of endogenous aSyn and the formation of LB-like inclusions. Its compatibility with the 96-well format also allows several compounds and concentrations to be compared in parallel under the same experimental conditions.

To determine whether selected protein turnover modulators affect the initiation of aggregation, primary mouse hippocampal neurons were first exposed to PBS or 70 nM mouse aSyn PFFs and co-treated with increasing concentrations of each compound or DMSO (Fig. 5A). The level of pS129-aSyn pathology was quantified by immunocytochemistry combined with high-content analysis (HCA; see Methods) 10 days after treatment (Figs. 5A–E, S6 and S7). The multiparametric staining strategy enabled quantitative discrimination of pS129-positive aSyn pathology within neuronal somata and MAP2-positive neurites, while simultaneously assessing neuronal survival (NeuN⁺/DAPI⁺), neuritic network integrity, and non-neuronal cell populations (DAPI⁺/NeuN⁻). Co-incubation of primary neurons with aSyn PFF and CGP-52411 or AS25224 resulted in a significant suppression of pS129 pathology at 10 µM or higher concentrations, consistent across all pS129 readouts, including total pS129 signal (Fig. 5C-D), as well as compartment-specific measurements within neuronal soma (Fig. S6A,C) and MAP2-positive neuritic processes (Fig. S6B,D). CI-994 also robustly reduced pS129 pathology in a concentration-dependent manner starting at 3 µM (Fig. 5B and 5E) in a comparable manner between the neuronal soma (Fig. S6E) and MAP2-positive neuritic processes (Fig. S6F).

**Fig. 5.**
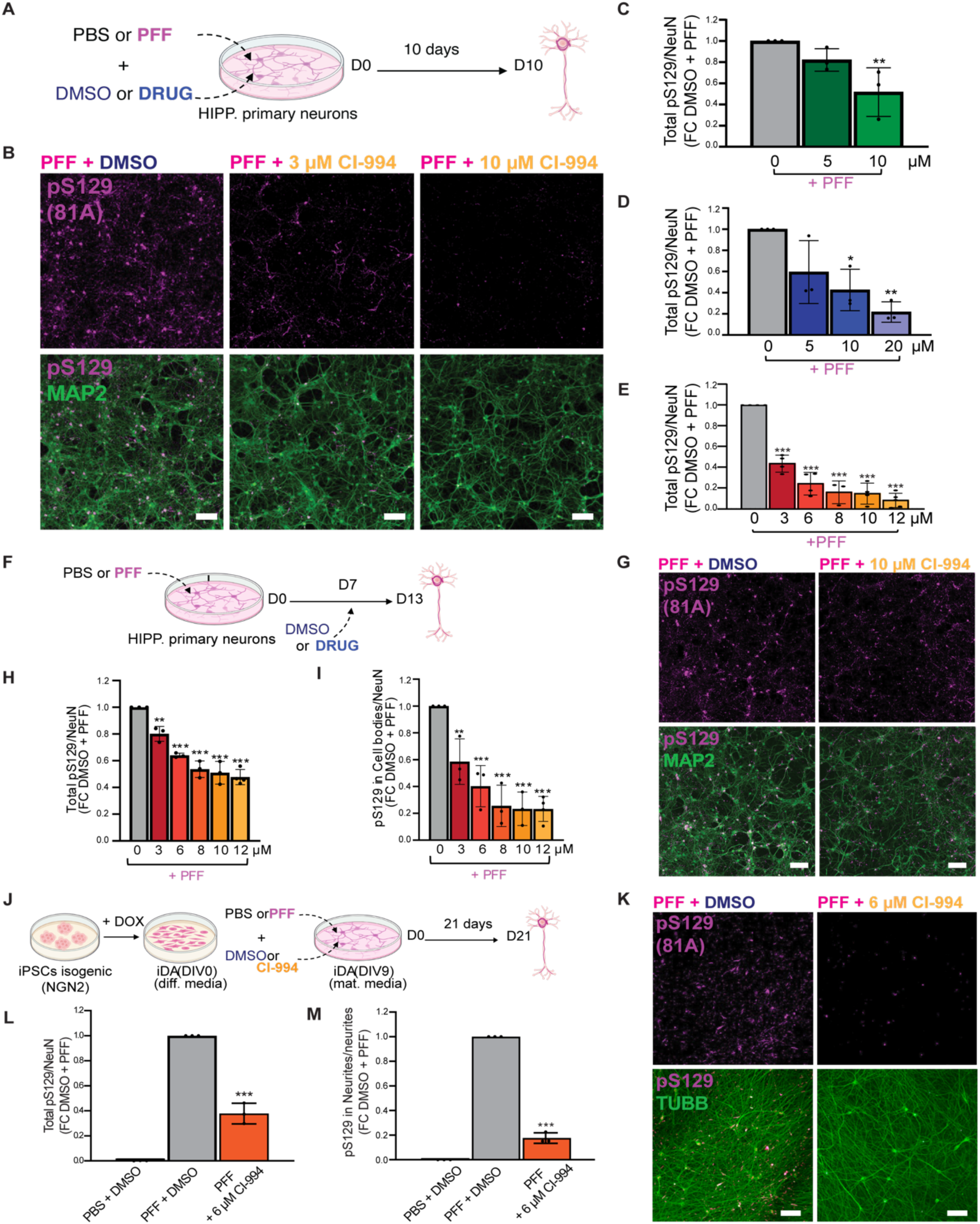
CI-994, AS-252424, and CGP-52411 reduce seeded pS129 aSyn pathology. **A.** Primary mouse hippocampal neurons were treated at DIV13 with PBS or mouse aSyn PFFs (70 nM) in the presence of DMSO or increasing concentrations of compounds and analyzed 10 days later (D10). **B.** Representative immunocytochemistry images of neurons treated with PFFs and co-treated with DMSO or CI-994 (3-10 µM). Pathological aSyn aggregates were detected using a phosphorylation-specific pS129 antibody (81A; magenta), and neuronal soma and neurites were detected with MAP2 (green). Merged images are shown below. Scale bar: 50 µm. **C-E**. Quantification of pS129-positive aSyn pathology by HCA. Total pS129 signal was normalized to PFF + DMSO controls following treatment with CGP-52411 (C), AS-252424 (D), or CI-994 (**E**). **F**. To assess effects on established pathology, neurons were exposed to PBS or mouse PFFs at DIV13 (D0). DMSO or compounds were added 7 days later (D7), and cultures were analyzed at D13. **G.** Representative images of PFF-treated neurons after delayed treatment with DMSO or CI-994 (10 µM). Pathological pS129-positive aggregates were detected with 81A (magenta), and neurons with MAP2 (green). Scale bar: 50 µm. **H–I.** Quantification of established pathology by HCA. Total pS129 signal (H) and somatic pS129 pathology within NeuN-positive cell bodies (**I**) were normalized to PFF + DMSO controls. **J.** Human iPSCs carrying a doxycycline-inducible NGN2 transgene were differentiated into dopaminergic neurons. At DIV9, neurons were treated with PBS or human aSyn PFFs together with DMSO or CI-994 (6 µM) and analyzed 21 days later. **K.** Representative images of iDA neurons treated with PFF + DMSO or PFF + CI-994. Pathological aggregates were detected using pS129 (81A; magenta), and neuronal processes were detected using βIII-tubulin (green). Scale bar: 50 µm. **L-M**. Quantification of total (**L**) and neuritic (**M**) pS129 pathology by HCA, expressed relative to PFF + DMSO controls. **C-E, H, I, L and M**. Data represent mean ± s.d.; N = 3 biological replicates with n = 3 technical replicates per biological replicate. *P < 0.05; **P < 0.01; ***P < 0.001.

Quantifications of the number of neurons, glial cells and neurites revealed only marginal changes induced by the different compounds, except for a modest decrease in the total number of neurons at the highest CI-994 concentrations (Fig. S5G-P and Fig. S8). These observations indicate that the suppression of pS129 pathology is not caused by cellular toxicity.

To independently validate the imaging-based findings, pS129 pathology was next assessed biochemically by Western Blot (WB) analysis of detergent-insoluble fractions, which are enriched in aggregated aSyn species (Fig. S6Q, S, U). Consistent with the HCA results, co-treatment with all compounds resulted in a pronounced reduction in pS129 immunoreactivity within the detergent-insoluble fraction (Fig. S6Q-V). The pS129 signal appeared as multiple bands, reflecting distinct phosphorylated aSyn species, a pattern characteristic of aggregated aSyn and comparable to that reported in human synucleinopathy brain tissue (*80*) and aggregation in the neuronal seeding model (*30*, *76–79*).

We next examined whether drugs influence endogenous aSyn levels, since these determine the amount of substrate available for seeded aggregation. Endogenous aSyn levels were quantified in PBS-treated neurons using the SYN-1 antibody to detect total endogenous aSyn at the single-cell level by immunocytochemistry combined with HCA (Fig. S6W). CI-994 induced a concentration-dependent reduction in endogenous aSyn levels (Fig. S6X), which was however, less pronounced than its effect on pS129 pathology. AS-252424 induced a small reduction in endogenous aSyn levels (∼20%) at the higher concentration tested (10 and 20 µM) (Fig. S6Y) and CGP-52411 did not induce detectable changes in endogenous aSyn levels (Fig. S6Z). Taken together, these results suggest that changes in endogenous aSyn levels do not account for the suppression of aggregation mediated by the compounds.

Having established that these drugs suppress seeded aggregation when co-applied with fibrils, we next asked whether they could also modulate aSyn pathology once aggregation had already been initiated. This question is particularly relevant in a therapeutic context, as disease symptoms are more likely to become visible only after pathological aggregates have begun to accumulate. To address this, we employed a delayed-treatment paradigm in which aSyn PFF were applied to primary neurons at day 13 in vitro (DIV13) to induce seeded aggregation, and compounds were added only after pathology had begun to develop. CI-994, AS-252424, or CGP-52411 were introduced 7 days after PFF exposure (D7) and maintained for an additional 6 days, allowing assessment of their ability to limit the accumulation or persistence of established aggregates (Fig. 5F). Using this delayed-treatment paradigm, pS129 pathology was quantified by HCA using the same multiparametric readouts described above. CI-994 reduced pS129 pathology in a concentration-dependent manner, with an approximately 40% reduction in total pathology at 6 µM (Fig.5G-H). Compartment-specific analyses revealed a stronger effect within neuronal soma than in neuritic processes (Fig.5I and S9A). Importantly, CI-994 did not induce detectable toxicity in this delayed-treatment paradigm, as neuronal counts, neuritic network density, and glial cell numbers remained comparable to PFF-treated DMSO controls (Fig. S9H-K). Under the same conditions, CI-994 also partially (∼40%) reduced endogenous aSyn levels (Fig. S9R-S). In contrast, delayed application of AS-252424 and CGP-52411 did not significantly reduce pS129 pathology at any of the concentrations tested (Fig. S9B-G). Total, somatic, and neuritic pS129 levels remained comparable to PFF-treated DMSO controls across all conditions. No compound-induced toxicity was detected in PBS-treated or PFF-treated cultures (Fig. S9L-Q), and endogenous aSyn levels remained unaltered (Fig. S9U-T). Together, these results indicate that only CI-994 suppresses aggregation both when present during the initiation phase and after aggregation has progressed.

We next asked whether this effect could arise from altered uptake or intracellular handling of aSyn PFF rather than from modulation of seeded aggregation itself. To do so, we used aSyn knockout (KO) neurons, which lack endogenous aSyn and therefore cannot support templated aggregation (*76*, *79*). aSyn KO neurons were exposed to aSyn PFF at DIV13 in the presence or absence of drugs, and intracellular fibril levels were quantified over time by WB (Fig. S9V-AA). In DMSO-treated cultures, internalized PFF were readily detected and underwent progressive intracellular processing and clearance, including characteristic proteolytic truncation into lower-molecular-weight C-terminal fragments, reflecting proteolytic processing of internalized seeds (Fig. S9V-X), as previously described (*79*). None of the compounds altered the amount of internalized PFF at early time points or their subsequent processing or degradation (Fig. S9Y-AA), suggesting that they act on the initiation phase of aggregation rather than on fibril uptake.

### CI-994 suppresses seeded aSyn aggregation in human iPSC-derived dopaminergic neurons

Following the initial screen in primary mouse neurons, CI-994 was selected as the most effective compound for validation in human iPSC-derived dopaminergic (iDA) neurons. The progressive loss of dopaminergic neurons is a central feature of PD, and this model reproduces the seeded aggregation of endogenous human aSyn and the progressive formation of LB-like pathology (*81*, *82*). However, pathology develops more slowly and at substantially lower levels than in primary mouse neurons (*82*), making the model less suitable for screening large numbers of compounds and concentrations. We therefore used iDA neurons for the validation of CI-994 in a human neuronal model of pS129-aSyn pathology.

At DIV9, iDA neurons were exposed to aSyn PFFs and treated with 6 µM CI-994. Pathology was assessed 21 days later, at DIV30 (Fig. 5J). Consistent with previous reports (*82*), seeded aggregates in iDA neurons developed more slowly than in primary mouse neurons and were initially and predominantly localized to neuritic compartments, with comparatively lower accumulation within neuronal soma at this time point (*83–86*). In these conditions, CI-994 treatment resulted in a robust reduction of pS129 pathology across all independent biological replicates (Fig. 5K, S10). Quantitative analysis revealed a pronounced decrease in total pS129 signal (Fig. 5L), with the strongest effect observed in neuritic processes (Fig. 5M), where pathological aggregates are preferentially localized in iDA neurons at day 21. Altogether, CI-994 emerged as the most promising compound due to its ability to suppress pS129 pathology in both primary mouse neurons and human iDA neurons, including under conditions where pathology is already established. While these results suggest that increased protein turnover elicited by CI-994 allows clearance of the pS129 pathology, the mechanistic link between the proteasome-related MCFT readout and the reduction in alpha-synuclein pathology remains to be established.

## Discussion

Here we describe a live-cell imaging approach to quantify global protein turnover in human neurons. The single-cell, quantitative nature of the readout obtained by a single microscopy snapshot makes this approach highly amenable to high-throughput screening starting from limited amounts of material. A single researcher can easily screen 2000 compounds per week in duplicates, a throughput vastly exceeding what is currently possible with proteomics-based approaches. Because it relies on a genetically encoded reporter, this approach is readily adaptable to different cell types and experimental contexts, making it well suited to screen for modulators of protein turnover in human pluripotent stem cell-derived differentiated cells in various developmental, disease, or perturbation conditions.

Our screen identified dozens of compounds that increase protein turnover, and we found the most promising molecules to increase the expression of ribosomal subunits and other translation-associated genes, providing a potential explanation for their ability to increase protein synthesis rate. This convergence on the translational machinery suggests that modulation of ribosome abundance or function may represent a common route to increase protein turnover in neurons. Whether the increase in ribosome subunits is caused by an increased production or decreased degradation of these proteins remains to be determined.

Surprisingly, even though CI-994 increased the protein levels of many ribosomal proteins, it did not increase their expression at the mRNA level. Such RNA–protein decoupling is well documented in the brain and reflects post-transcriptional regulation, including differences in translation, trafficking and protein turnover (*11*, *24*, *87*, *88*). We did not observe consistent changes in the expression of mRNAs and proteins involved in proteasomal or autophagic protein degradation. This suggests that the increase in protein degradation we measured is driven by changes in the activity of the degradation machinery, which can be mediated by assembly and accessory regulation such as regulator/cap binding, chaperones, substrate ubiquitination, and/or proteasome PTMs without coordinated transcriptional changes (*89*, *90*). Altered degradation activity with unchanged proteasome levels is also consistent with the documented lack of correlation between proteasome abundance and activity in quiescent cells (*91*), which in some neuronal types could be explained by the maintenance of substantial spare proteasomal capacity, with only 20 % of 26S proteasomes actively engaged in substrate processing (*92*).

How the three selected compounds modulate ribosomal protein expression and protein turnover in human neurons remains unclear. AS-252424 is a selective PI3Kγ inhibitor (*93*), while CGP-52411 is an EGFR kinase inhibitor (*94*). Both act on the PI3K/Akt/mTORC1 signaling axis, a central regulator of growth and protein synthesis (*43*, *95*). EGFR functions also upstream of Ras/Erk, which can feed into mTORC1 and further support anabolic programs (*43*, *96*). PI3K signaling also interfaces with mTORC2, including reported effects on mTORC2–ribosome association (*97*, *98*). EGFR/PI3K inhibition would be expected to dampen mTORC1-driven protein synthesis, with the potential secondary engagement of catabolic programs through reduced mTORC1/C2 activity. However, in senescent cells mTORC1 regulation can become uncoupled from normal nutrient and growth factor sensing, resulting in persistent mTORC1 activity despite upstream deprivation (*99*). Although our system differs from senescence, this observation illustrates how altered pathway coupling in nondividing cells may reshape proteostasis in ways not readily inferred from proliferating cellular models. Consistent with this idea, we did not observe downregulation of genes or proteins upregulated by the mTOR pathway following CGP-52411 or AS-252424 treatment, even though both compounds upregulated ribosomal mRNA and protein levels. While CI-994 is a class I HDAC inhibitor (*72*), the limited changes observed at the proteome level despite broad transcriptional reprogramming suggest that the effects on protein turnover are primarily controlled at the post-transcriptional and protein degradation levels. Recent evidence links CI-994 to Wnt/β-catenin signaling (*100*), which activates mTORC1 and promotes translation (*101*, *102*). HDAC inhibition has also been reported to increase acetylation of 20S proteasome subunits (*103*), enhancing proteolytic capacity, and in some contexts to activate TFEB-dependent lysosomal biogenesis (*104*). Interestingly, in neurons both class I HDAC inhibitors, including CI-994, and perturbation of the PI3K/mTOR pathway have been linked to activity-and plasticity-related phenotypes (*73*, *100*, *105–107*). Memory consolidation depends on regulated protein homeostasis, including de novo protein synthesis, mTORC1 signaling, and efficient UPS function (*107–110*). Furthermore, neuronal activity and plasticity are accompanied by enhanced remodeling of protein turnover programs and changes in synaptic protein stability (*4*, *111–113*). Although we did not investigate this connection, these observations raise the possibility that these apparently distinct mechanisms intersect at the level of neuronal proteostasis. Nevertheless, it is plausible that these compounds exert broader, off-target activities beyond their canonical targets, as effects independent of their primary mechanisms of action have been reported (*100*, *114*, *115*). While all three compounds induced transcriptional changes, alterations of the neuronal proteome were relatively limited, suggesting that cell identity was not significantly altered after acute drug treatment. CI-994 suppressed pathological features in all mouse and human in vitro models of PD that we tested, consistent with its reported neuroprotective effects in other preclinical settings, including promotion of functional recovery after traumatic brain injury and reduction of tau phosphorylation in cellular AD models (*100*, *105*). Altogether, these findings position CI-994 as a compound of potential translational relevance and motivate further work to disentangle how CI-994 reshapes neuronal stress and proteostasis pathways.

In summary, our work establishes a new platform that considerably facilitates and increases the throughput of global protein synthesis and degradation rate measurements, allowing for the rapid discovery of biomedically-relevant compounds increasing protein turnover. This approach can be easily implemented in any cell type derived from our established pluripotent stem cell line, or in which the MCFT can be stably expressed, thereby holding broad potential for screening protein turnover modulators across a wide range of physiological and pathological in vitro cellular models.

## Limitations of this study

One important limitation of our study is the use of in vitro, iNGN-generated neurons to measure the impact of small molecules on protein turnover. We cannot exclude that more mature neurons and/or different neuronal subtypes found in the human brain may respond differently to these compounds. However, the efficacy of CI-994 in reducing pathological features of PD both in mouse primary neurons and iPSC-derived dopaminergic neurons suggests its broad applicability to increase protein turnover in neurons. We also assume that the 24-h post-perturbation time point approximates steady state conditions, enabling protein turnover to be inferred from a single snapshot of the timer. However, we cannot rule out that the activity of some pathways or the level of specific proteins is still dynamically changing at this time point, which could influence the interpretation of turnover measurements.

## Acknowledgments

Microscopy was performed using the resources of the EPFL Bioimaging and Optics Core Facility (EPFL-BIOP). Mass spectrometry was performed at the Proteomics Core Facility (EPFL-PCF), with special thanks to Maria Pavlou and Florence Armand. Drug screening was performed at the EPFL Biomolecular Screening Facility. We would like to express our appreciation to Gerardo Turcatti, Fabien Kuttler and Julien Chapalay. RNA sequencing was performed at EPFL Gene Expression Core Facility. We would like to kindly thank Benjamin Martin for adjusting the Python scripts to high throughput needs of drug screening, and Giovanni d’Angelo for critical reading of the manuscript. Icons in Fig. 2A are licensed under CC-BY 3.0 or 4.0 Unported https://creativecommons.org/licenses/by/4.0/. Icons in Fig. 5 were created in Biorender.com MELLIER, A. (2026) https://BioRender.com/e51lpnt.

## Funding

EPFL core funding

Synapsis Foundation Switzerland (grant no. 2021-PI03)

Fondation Bru

## Author contributions

Conceptualization, JD, ALM, DMS

Methodology, JD, ALM, DMS

Validation, JD, ALM

Formal Analysis, JD, ALM, AT

Investigation, JD, ALM, AT, YJ

Data Curation, JD, ALM, AT

Writing – Original Draft, JD, ALM, DMS

Visualization, JD, ALM, AT

Supervision, ALM, DMS

Funding Acquisition, ALM, DMS

## Data and materials availability

Plasmids and cell lines are available from the lead contact upon request. LFQ Proteomics raw data have been deposited in the PRIDE database (PXD077075). RNA-seq data have been deposited to the GEO database with accession number GSE328822. Imaging data or other data reported in this paper are available from the lead contact upon request. Any additional information required to reanalyze the data reported in this paper is available from the lead contact upon request.

## Code availability

No new code was developed specifically for the analyses presented here. All software tools and previously developed code are stated and cited in the Methods.

## Materials and Methods

### Ethical statement

All animal procedures were performed in compliance with institutional and national regulations and were approved by the Swiss Federal Veterinary Office (authorization VD4029). All experiments involving hESCs and iDA (approval number 2025-00664) were approved by the Canton of Vaud Ethics committee on human research (https://www.cer-vd.ch).

### Cell lines used in this study

The generation of human embryonic stem cells (hESCs) expressing the tandem fluorescent timer (FT) and/or neurogenin 1 and neurogenin 2 (NGN1/2) with the TRE promoter used in this study has been previously described in (*42*). For aSyn seeding experiments, NGN2-inducible iPSCs (AIW002-02) were obtained from Early Drug Discovery Unit, McGill University. All cell lines were routinely checked for mycoplasma and tested negative.

### Cell culture and maintenance

hESCs were maintained as previously described (*42*). Briefly, hESCs were routinely cultured as colonies in mTeSR Plus (STEMCELL Technologies, 100-0276) in Corning Matrigel hESC-qualified matrix-coated cultureware (Corning, 354277) at 37 °C, 5% CO_2_. Cells were passaged once or twice per week using the enzyme-free passaging reagent ReLeSR (STEMCELL Technologies, 05872). For single cell dissociation, hESCs were harvested using Accutase (Innovative Cell Technology, AT104). After 5 min incubation at 37 °C, DMEM/F12 containing 15 mM HEPES (STEMCELL Technologies, 36254) was added, and the cell suspension was centrifuged 5 min at 300 x g. The supernatant was discarded, and cells were resuspended in fresh mTeSR Plus with 10 µM ROCK Inhibitor (Y-27632) (MilliporeSigma, SCM075).

### Neuronal induction

hESCs were induced to neurons (iNGNs) as previously described (*42*). Briefly, hESCs were cultured for at least two passages prior to the induction. hESCs were harvested as single cells using Accutase (Innovative Cell Technology, AT104) and plated at 5 x 10^4^ cells/cm^2^ in mTeSR Plus medium (STEMCELL Technologies, 100-0276) supplemented with 10 µM ROCK inhibitor (MilliporeSigma, SCM075) in cultureware coated with 15 µg/mL Poly-L-Ornithine (PLO, Sigma-Aldrich, P4957) and 10 µg/mL laminin (Sigma-Aldrich, L2020), resuspended in PBS with Mg^2+^ and Ca^2+^. For the next four days, the mTeSR Plus medium was supplemented with 1 µg/mL doxycycline (Sigma-Aldrich, D9891) and changed daily. For the last 24 h, the medium was additionally supplemented with 5 µM cytosine β-D-arabinofuranoside hydrochloride (Sigma-Aldrich, C6645). Afterwards, iNGNs were maintained in BrainPhys™ Neuronal Medium (STEMCELL Technologies, 05790) or BrainPhys™ Imaging Optimized Medium (STEMCELL Technologies, 05796) supplemented with a final concentration of 1x N2 Supplement-A (STEMCELL Technologies, 07152), 1x NeuroCult™ SM1 Neuronal Supplement (STEMCELL Technologies, 05711), 20 ng/mL BDNF (STEMCELL Technologies, 78005), 20 ng/mL GDNF (STEMCELL Technologies, 78058), 200 nM ascorbic acid (STEMCELL Technologies, 72132) and 1 mM dibutyryl-cAMP (STEMCELL Technologies, 73882). Half of the medium was changed twice a week.

### Neuronal 2D long-term differentiation

#### Differentiation into neural progenitor cells (NPCs)

hESCs were maintained in culture two weeks prior to the start of differentiation, followed by passaging as single cells as described above and seeded at 2.5 x 10^5^ cells/cm^2^ in STEMdiff Neural Induction Medium with 1x STEMdiff SMADi Neural Induction Supplement (STEMCELL Technologies, 08581) with 10 µM ROCK Inhibitor (Y-27632) (MilliporeSigma, SCM075) on Corning Matrigel hESC-qualified matrix-coated cultureware (Corning, 354277), according to the manufacturer’s protocol and as described earlier in (*42*). Medium change was performed daily for three weeks. NPCs were passaged using Accutase (Innovative Cell Technology, AT104) once a week.

#### Differentiation into neurons

NPCs were differentiated into neurons as described earlier with minimal modifications (*66*) and according to the manufacturer’s protocol. NPCs were passaged using Accutase (Innovative Cell Technology, AT104) and plated at 5 x 10^4^ cells/cm^2^ in STEMdiff Neural Induction Medium in cultureware coated with 15 µg/mL Poly-L-Ornithine (PLO, Sigma-Aldrich, P4957) and 10 µg/mL laminin (Sigma-Aldrich, L2020), resuspended in PBS with Mg^2+^ and Ca^2+^. The next day, half of the medium was changed to BrainPhys™ Neuronal Medium (STEMCELL Technologies, 05790) or BrainPhys™ Imaging Optimized Medium (STEMCELL Technologies, 05796) supplemented with a final concentration of 1x N2 Supplement-A (STEMCELL Technologies, 07152), 1x NeuroCult™ SM1 Neuronal Supplement (STEMCELL Technologies, 05711), 20 ng/mL BDNF (STEMCELL Technologies, 78005), 20 ng/mL GDNF (STEMCELL Technologies, 78058), 200 nM ascorbic acid (STEMCELL Technologies, 72132) and 1 mM dibutyryl-cAMP (STEMCELL Technologies, 73882). Half-medium change was performed every two to three days until imaging.

### Live cell imaging

Imaging was performed as previously described (*42*). 96-well black imaging plates (PerkinElmer, 6055302) were coated with Corning Matrigel hESC-qualified matrix-coated cultureware (Corning, 354277). One day prior to imaging, cells were seeded as single cells as described above. A snapshot image was taken with the Operetta CLS microscope (PerkinElmer), 20× objective (Air immersion, NA 0.8), at 37 °C, 5% CO_2_. For the green and red channels, the following filters were used, respectively: Ex: BP 435-460, 460-490, Em: HC 500-550 and Ex: BP 490-515, 530-560, Em: HC 570-650.

### Immunofluorescence for validation of differentiation

Immunofluorescence was performed as previously described (*42*) with minimal modifications. 0.1 M PHEM buffer (Electron Microscopy Sciences, 11162) was added to the neuronal medium in equal amounts. Cells were washed twice with 0.1 M PHEM buffer (Electron Microscopy Sciences, 11162) and fixed with 4 % formaldehyde (Thermo Fisher Scientific, 28906), 0.25 % glutaraldehyde (Electron Microscopy Sciences, 16220) in 0.1 M PHEM buffer for 15 min at room temperature. Cells were washed twice with PBS (Bioconcept, 9872L), permeabilized with 0.1 % Triton X-100 (BioChemica, UN3082) in PBS for 20 min and blocked with 1 % BSA (Sigma-Aldrich, A7906) in PBS for 30 min. Cells were incubated overnight at 4 °C with anti-Ki67 antibody (1:100, BD Biosciences, 550609), anti-MAP2 antibody (1:500, Sigma-Aldrich AB5622), or anti-TUBB antibody (1:2000, eBioscience 14-4510). The next day, cells were washed twice with PBS (Bioconcept, 9872L) and incubated with an anti-mouse secondary antibody conjugated to AlexaFluor647 (1:1000, LifeTechnologies, A31571) or an anti-rabbit secondary antibody conjugated to AlexaFluor647 (1:1000, LifeTechnologies, A21443) for 1 h at room temperature. Cells were washed twice with PBS (Bioconcept, 9872L) and mounted with VECTASHIELD® HardSet™ Antifade Mounting Medium with DAPI (Vector, H-1500-10). Cells were imaged as described above with the far-red filters: Ex: BP 615-645, BP 650-675, Em: HC 655-760.

### Image processing

Image processing was performed as previously described (*42*). Briefly, all images were background-corrected to account for uneven illumination and auto-fluorescence of the medium, while the dark field signal was neglected as it was negligible. The illumination pattern was obtained by imaging a well containing only medium under identical exposure settings and normalizing the image to its mean intensity. Raw images were then divided by this reference to generate flat field corrected images. Auto-fluorescence was estimated for each frame individually by applying a thresholding method to the corrected images, creating a binary mask that excluded foreground fluorescence signals. The threshold was set to the fifth percentile of the pixel intensity distribution within segmented cells, and the mask was expanded by erosion. The mean or peak pixel intensity was measured from the unmasked regions and used for background subtraction for each individual cell. Cell nuclei were segmented in the green fluorescence channel using CellPose 2.0 with the built-in “nuclei” model in Python, generating masks that were subsequently used to extract green and red fluorescence intensities for each cell (*116*). Microscopy Figures were created using the microfilm package in Python (*117*).

### Quantification of MCFT half-life

The half-life (t_1/2_) of the MCFT was calculated as previously described (*42*). Briefly, the decay rate (k) was calculated from the following equation:

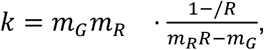

where m_G_ - maturation rate of sfGFP; m_R_ - maturation rate of mOrange2; R - green to red fluorescence ratio based on the integrated fluorescence intensity. Maturation rates of sfGFP and mOrange2 were taken from (*42*). Note that *k* = *k_de_*_g*radation*_+*k_dilution_*.

Based on k, the t_1/2_ was computed, filtering for k>0:

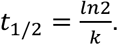

### Drug Screening

Compounds from the EPFL’s Biomolecular Screening Facility (BSF) in-house collections of 257 Kinase Inhibitors (Supplementary File S1), 4360 molecules from in-house Drug Repurposing Library (selected from the Broad Institute Drug Repurposing Library, Supplementary File S2), and 1280 molecules from Prestwick Chemical Library (Prestwick Chemicals) were used for the drug screening. Compounds were provided at 10 mM stock solutions in DMSO, they were flushed with Argon and stored at –20 °C under dry air, protected from light. Chemical integrity of the libraries was controlled by HPLC-MS. Drugs were spotted on a PhenoPlate™ 384-well plate (Revvity, 6057302) using an acoustic liquid handler (Beckman Coulter Echo 655). Plates were stored at – 20 °C.

### Screening window coefficient

To determine the quality of the assay, the screening window coefficient (Z’-factor) was calculated as follows:

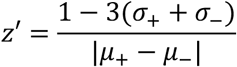

where σ is the standard deviation and µ is the mean of fluorescence intensity of the positive and negative controls, respectively (*67*). As a validation of the assay, z’-value was assessed on a z’-plate composed of 192 replicates of DMSO (negative control) and 192 replicates of 20 µM final cycloheximide (CHX; inverse (positive) control). The quality of the assay was deemed as sufficient if the obtained z’ was above 0.5. FT-iNGNs were differentiated as described above, harvested using Accutase (5 min, 37 °C) resuspended in DMEM/F12 with 15 mM HEPES (STEMCELL Technologies, 36254), and then centrifuged 5 min at 400 x g. The cell pellet was resuspended in Brain Phys™ Imaging Optimized Medium (STEMCELL Technologies, 05796) supplemented as described above to 7.5 x 10^5^ cells/mL. Laminin (Sigma-Aldrich, L2020) at the final concentration of 10 µg/mL was added to the cell suspension to ensure cell attachment. Automatic cell seeding into 384-well imaging plates was performed using a Multidrop Combi (Thermo Fisher) dispenser at medium speed, dispensing 20 µL of cell suspension per well. After 24 h incubation at 37 °C with 5% CO_2_, fluorescence snapshots of the green and red channels were acquired on the Operetta high-throughput fluorescence microscope (Perkin Elmer) equipped with a 20x/0.8 NA objective, at 37 °C with 5% CO_2_. For each well, 9 fields were acquired. The primary readout considered was the mean intensity ratio of green versus red fluorescence channels and the data were processed as explained in the Image processing section.

### Primary screening

Drugs were tested at a final concentration of 10 µM, in biological duplicates (on separate plates). Each plate contained controls used for per-plate and per-batch normalization and z’-factor determination, with two columns per plate: DMSO and 20 µM CHX. The plates were spotted, and cells were seeded as described above. The list of compounds from the in-house libraries are provided in Supplementary files S1 and S2.

#### Primary screening data curation

The Laboratory Information Management System (LIMS) developed by EPFL’s BSF was used for data processing and statistical validation of hits. Only plates with a z’-factor above 0.25 were considered for further analysis as calculated based on the control wells from each plate. Data were normalized by assigning 0 to the mean G/R ratio of DMSO control and 1 to the G/R ratio of CHX. A normalized average score was calculated for each compound based on the duplicate. A compound was considered a validated hit if its score was different from the mean of the DMSO control + 3 standard deviations. Cell viability was assessed in parallel based on the segmented cell count, as a proxy for the toxicity of screened compounds.

### Secondary screening

For the secondary screening, the data from the primary screen were manually curated by removing hits resulting from imaging artifacts, compounds decreasing the cell number by more than 2.5-fold as compared to the DMSO control, and structural analogs. The drugs sharing the same mechanism of action were limited to no more than three. The selected 47 drugs were tested in dose-response with ten final concentrations ranging from 3 nM to 40 µM in duplicates, using the same protocol as for the primary screen. The half maximal effective concentrations (EC50) were determined for each compound by fitting the dose-response to a sigmoidal function.

#### Selected compounds

For the follow-up experiments, the selected compounds CGP-52411 (HY-103442), AS-252424 (HY-13532), and CI-994 (HY-50934), were acquired from MedChemExpress, resuspended in 99.9% DMSO (Sigma Aldrich, 900645) at 50 mM and stored in aliquots at-80 °C.

### Label Free Quantitative (LFQ) Proteomics

LFQ Proteomics was performed as described previously with minor modifications to adjust to neuronal culture (*42*). Proteomics experiments were performed by the Proteomics Core Facility at EPFL. The experiment was performed in N = 3 biological replicates.

#### Cell culture

Upon thawing, hESCs were maintained for two weeks as described above. iNGNs were induced in a 12-well plate as described above, adjusting the volumes and cell number accordingly to the well size. Directly after the end of induction, cells were treated with 20 µM CGP-52411 (HY-103442), 20 µM AS-252424 (HY-13532), 5 µM CI-994 (HY-50934) or a volume of DMSO corresponding to the highest drug concentration resuspended in BrainPhys™ Neuronal Medium (STEMCELL Technologies, 05790) completed with all supplements as described earlier. iNGNs were incubated for 24 h at 37 °C with 5% CO_2_.

#### Sample collection

Cell culture dishes were placed on ice, the medium was aspirated and cells were washed twice with ice-cold PBS without Mg^2+^ and Ca^2+^. Cells were then scraped in 50 µL of lysis buffer (100 mM Tris buffer pH 8 (PanReac AppliChem, A4577), 2% SDS (PanReac AppliChem, A3942), 1x Halt Protease Inhibitor (LifeTechnologies, 78441)) and collected in protein low-binding tubes. Benzonase nuclease (Sigma-Aldrich, 70746-3) was added to each tube and incubated for 15 min at room temperature. The lysates were boiled at 90 °C for 10 min and centrifuged at 20,000 x g for 10 min at 4 °C. The protein extracts were transferred to a new tube without disturbing the pellet and snap-frozen in liquid nitrogen. They were stored at-80 °C until further use. The protein extract concentration was measured using the Pierce™ BCA Protein Assay Kits (Thermofisher, 23225).

#### Sample preparation

Protein samples (20 µg) were processed using a filter-aided sample preparation (FASP) workflow with slight adaptations (*118*). Briefly, proteins were loaded onto pre-washed and equilibrated Microcon®-30K centrifugal filter units (Merck AG, Zug, Switzerland) and centrifuged at 9,400 × g at 20 °C for 30 minutes, or until complete dryness was achieved. All subsequent centrifugation steps were carried out under identical conditions. The retained proteins were washed twice with 200 µL of urea buffer (8 M urea, 100 mM Tris-HCl, pH 8.0). Reduction was performed by applying 100 µL of 10 mM Tris(2-carboxyethyl)phosphine (TCEP) prepared in urea solution, followed by incubation for 60 minutes at 37 °C with gentle agitation under light-protected conditions. The reduction solution was removed by centrifugation and washed twice with 200 μL urea solution. Alkylation was then carried out by adding 100 µL of 40 mM chloroacetamide (CAA) to the urea buffer and incubating for 45 minutes at 37 °C with gentle shaking in the dark. The alkylation solution was removed by centrifugation, followed by two additional washes with the urea buffer. Subsequently, two conditioning washes were performed using 200 µL of 5 mM Tris-HCl (pH 8.0) to prepare the samples for enzymatic digestion. Proteins were digested overnight at 37 °C by adding 100 µL of a protease mixture containing endoproteinase Lys-C and trypsin (Trypsin Gold) at an enzyme-to-protein ratio of 1:50 (w/w), prepared in 5 mM Tris-HCl supplemented with 10 mM CaCl₂. Peptides were recovered by centrifugation and further eluted with two sequential additions of 50 µL 4% trifluoroacetic acid. The resulting peptide mixtures were desalted using SDB-RPS StageTips (*119*) and dried by vacuum centrifugation prior to LC–MS/MS analysis.

#### Mass Spectrometry

Dried peptide samples were resuspended in 2% acetonitrile (Biosolve) containing 0.1% formic acid. Peptide separation was performed using a Dionex Ultimate 3000 RSLC nano-UPLC system (Thermo Fisher Scientific) coupled online to an Orbitrap Ascend mass spectrometer (Thermo Fisher Scientific). Samples were first loaded onto a trapping column (Acclaim PepMap C18, 3 µm, 100 Å, 2 cm × 75 µm internal diameter (ID)) for desalting and concentration. Analytical separation was achieved on a 50 cm capillary column (75 µm ID), packed in-house with ReproSil-Pur C18-AQ 1.9 µm silica particles (Dr. Maisch), using a flow rate of 250 nL/min and a 150-minute biphasic gradient. Data acquisition was performed in Top Speed data-dependent acquisition (DDA) mode with a cycle time of 2 seconds. Full MS scans were recorded at a resolution of 60,000 (at m/z 200). The most intense precursor ions were selected for fragmentation by higher-energy collisional dissociation (HCD) using a normalized collision energy (NCE) of 30% and an isolation window of 1.4 m/z. Fragment ion spectra were acquired at a resolution of 15,000 (at m/z 200), and previously fragmented ions were dynamically excluded for 20 seconds.

#### Data analysis

Raw mass spectrometry data were analyzed using MaxQuant (version 2.4.4.0) (*120*) against a protein database containing 104557 entries (LR2024_01). Carbamidomethylation of cysteine residues was specified as a fixed modification, while methionine oxidation, phosphorylation (serine, threonine, tyrosine), protein N-terminal acetylation, and glutamine conversion to pyroglutamate were included as variable modifications. Up to two missed cleavages were permitted, and the “match between runs” feature was enabled. Protein identification required at least two peptides, and the false discovery rate (FDR) was controlled at 0.01 at both peptide and protein levels. Label-free quantification (LFQ) was carried out using the MaxLFQ algorithm with default parameters (*121*).

#### Statistical Analysis

Downstream statistical analyses were conducted using Perseus (version 1.6.12.0) (*122*). Reverse hits, contaminants, and proteins identified only by site were excluded. Protein groups with at least three valid intensity values in at least one experimental condition were retained. Missing values were imputed using a normal distribution (width = 0.4, downshift = 1.8 standard deviations). On the graphs, each point is the average over biological triplicate of the protein level (P) fold change compared to the vehicle treatment (DMSO). Red point marks significant protein groups above the curved blue line based on the differential protein abundance between conditions using a two-sample t-test with permutation-based FDR correction (250 permutations, FDR = 0.05, S0 = 0.1). Group-level analysis of predefined protein sets was performed based on the lists of ribosomal (RPS, RPL) and proteasomal proteins that were taken from (*123*)(retrieved January 2026). Autophagy (GO:0006914), mTOR signaling (GO0031929), positive mTOR regulation (GO0032008), and negative mTOR regulation (GO0032007) were taken from Gene Ontology enrichment list on QuickGO (*124*). For each set, protein values were compared with those of all remaining quantified proteins using a two-sided Wilcoxon-Mann-Whitney test (*125*), and P values were adjusted across tested sets by the Benjamini-Hochberg method (FDR = 2%) (*126*).

### RNA sequencing

#### Cell culture

Cells were cultured the same way as for LFQ proteomics. RNA sequencing was performed at EPFL Gene Expression Core Facility. The experiment was performed in N = 2 biological replicates.

#### Sample collection

After 24 h incubation, cells were washed twice with ice-cold PBS without Mg^2+^ and Ca^2+^, scraped manually, subsequently transferred to a DNA LoBind Tube (Eppendorf, 022431021) and centrifuged at 4 °C at 1000 x g. The supernatant was removed and cell pellets were snap-frozen in liquid nitrogen and stored at-80°C until further processing.

#### Sample preparation

RNA were extracted using RNeasy Plus Micro Kit (Qiagen #74034) following manufacturer’s instructions with an additional step of on-column DNAse digestion (Qiagen, #79254). Sequencing libraries were prepared on 500 ng RNA by the EPFL Gene Expression Core Facility using the NEBNext Ultra II Directional RNA Library Prep with Ribodepletion Kit for Illumina (NEB, #E7760S).

#### Sequencing

Libraries were sequenced on an Aviti24 sequencer (Element Biosciences) using 75-nucleotide length paired-end sequencing.

#### Data analysis

RNA-Seq libraries were mapped to the hg38 human genome using STAR (*127*). Exon coordinates were gathered from GENCODE annotation from v38 of the UCSC Genome Browser. Gene counts were then determined using the featureCounts function (*128*) from the R package Rsubread. Differential enrichment analysis as well as normalization of results using the lfcShrink function with type =”normal” were performed using DeSeq2 (*129*). Gene ontology enrichment analysis was performed with the web tool, ShinyGO 0.85.1 (*130*), using Curated.Reactome as pathway database. The top 15 pathways were selected by fold enrichment, filtered by FDR < 0.05 and displayed ranked by FDR.

Cytosolic ribosomal proteins (WP477), proteasome degradation (WP183), PI3K pathway (WP4172), EGFR pathway (WP4806)-related genes were taken from WikiPathways (*131*). Genes related to autophagy (GO:0006914), mTOR (GO:0031929) and neurogenesis (GO:0022008) were taken from the Gene Ontology enrichment list on QuickGO (*124*).

### Expression, purification, and fibrillization of mouse aSyn

Recombinant wild-type mouse aSyn was produced in *Escherichia coli* using a pT7-7 expression system. Following bacterial expression, aSyn was purified through sequential anion-exchange chromatography and reverse-phase high-performance liquid chromatography, as described previously (*132*, *133*). Purified monomeric protein was diluted in PBS (Thermo Fisher Scientific, #10010015) at pH 7.5 to a final concentration of 170-200 µM, passed through a 100-kDa molecular weight cutoff filter (Millipore, MRCF0R100) to remove pre-existing aggregates, and incubated at 37 °C under continuous orbital agitation (1,000 rpm) for 5 days to promote fibril assembly (*134*). The formation of β-sheet-rich fibrillar structures was verified by thioflavin T (Sigma-Aldrich, 596200) fluorescence measurements (*134*). To generate preformed fibril (PFF) seeds suitable for neuronal seeding assays, fibrils were fragmented by controlled probe sonication on ice (total sonication time 20 s, 20% amplitude, 1 s on/1 s off), yielding short fibrillar species with an average length of approximately 50-100 nm and minimal release of monomeric aSyn (*76*). Sonicated mouse aSyn PFF were characterized by Coomassie staining and transmission electron microscopy (*134*), aliquoted, snap-frozen in liquid nitrogen, and stored at −80 °C until use.

### Primary hippocampal neuron cultures

Primary hippocampal neurons were generated from postnatal day 0-1 (P0-P1) mouse pups derived from either wild-type (WT) C57BL/6J or aSyn knockout (aSyn KO; OLA) backgrounds. Timed-pregnant C57BL/6Jrj females were obtained from Janvier Labs (France) and C57BL/6JOlaHsd females from Envigo (France). Pregnant dams at embryonic day 12 were received at the EPFL animal facility and kept under controlled housing conditions through delivery. Neonatal pups were euthanized by decapitation immediately prior to tissue collection, and all experiments were conducted ex vivo. In accordance with the 3Rs principles, dams were reassigned to the EPFL Organ/Tissue Sharing Program (Optimice) whenever possible. All animal procedures were performed in compliance with institutional and national regulations and were approved by the Swiss Federal Veterinary Office (authorization VD4029). Under sterile conditions, hippocampi were rapidly dissected in ice-cold Hank’s Balanced Salt Solution (HBSS, Thermo Fisher Scientific 314025050) and enzymatically dissociated using papain (20 U/mL) (Sigma-Aldrich, P3125) for 30 min at 37 °C. Enzymatic digestion was terminated by the addition of a protease inhibitor solution, followed by gentle mechanical trituration to obtain a single-cell suspension. Cells were collected by centrifugation, resuspended in Neurobasal medium (Thermo Fisher Scientific, 21103049), supplemented with B27 (Thermo Fisher Scientific, 17504044), L-glutamine, and penicillin/streptomycin, and plated onto poly-L-lysine (Biotechne, 3438-100) - coated black, clear-bottom 96-well plates at a density of 200,000 cells/mL (200 µL per well). Cultures were maintained at 37 °C in a humidified incubator with 5% CO₂ and were left undisturbed except for experimental treatments.

### Differentiation of NGN2-iPSCs into dopaminergic neuronal cultures (iDA)

Human NGN2-inducible iPSCs (AIW002-02; Early Drug Discovery Unit, McGill University) were differentiated into induced dopaminergic (iDA) neuronal cultures using an optimized protocol previously established (*82*, *135*). Briefly, NGN2-expressing iPSCs were first differentiated into induced neurons. Cells were plated one day prior to induction in mTeSR Plus medium (STEMCELL Technologies, 100-0276) supplemented with the ROCK inhibitor Y-27632 (Selleckchem, S1049). On the day of induction (DIV0), NGN2 expression was initiated by replacing the medium with a defined differentiation medium containing doxycycline, neurotrophic factors, and laminin. Medium was refreshed on DIV1. At DIV2, cells were dissociated with Accutase and replated onto poly-L-ornithine-and laminin-coated plates or coverslips in Neurobasal medium supplemented with N2, B27, GlutaMax, neurotrophic factors, laminin (Thermo Fisher Scientific, 23017-015), and doxycycline. From DIV3 onward, cultures were transitioned to a midbrain dopaminergic differentiation medium (STEMdiff Midbrain Neuron Differentiation Kit, STEMCELL Technologies, 100-0038) supplemented with doxycycline and Sonic Hedgehog (STEMCELL Technologies, 78065). Medium was refreshed at DIV6. At DIV9, cultures were switched to maturation medium from the STEMdiff Midbrain Neuron Maturation Kit (STEMCELL Technologies, 100-0041). From this stage onward, half of the medium was replaced weekly with fresh maturation medium. iDA cultures were maintained under these conditions until experimental use.

### Treatment of primary hippocampal neurons or iDA with mouse aSyn PFF and small-molecule modulators

For aggregation assays, primary hippocampal neurons or human iDA neurons were exposed to mouse aSyn PFF using two complementary experimental paradigms. Mouse aSyn PFF aliquots were thawed at room temperature immediately prior to use and diluted in conditioned Neurobasal medium collected from the corresponding neuronal cultures. PFF were applied at a final concentration of 70 nM in primary hippocampal neurons and 500 nM in human iDA neurons.

In the co-treatment paradigm, aSyn PFF were applied concurrently with small-molecule modulators to assess their effects on the initiation and early accumulation of seeded aggregation. In primary mouse hippocampal neurons, aSyn PFF were added at day in vitro 13 (DIV13; D0) together with the indicated concentrations of CI-994, AS-252424, or CGP-52411. Neurons were maintained under these conditions for 10 days, and aSyn pathology and cellular parameters were assessed at D10 post-treatment (DIV23). In parallel experiments using human induced dopaminergic (iDA) neurons, aSyn PFF were added at DIV9 together with 6 µM CI-994. Cultures were maintained for 21 days following treatment, and analyses were performed at DIV30 (D21). This extended treatment window was selected to accommodate the slower maturation and aggregation kinetics of human neurons. For both mouse and human neuronal cultures, compounds were prepared as described earlier, diluted into the corresponding neuronal culture medium to generate intermediate working solutions, and then added to the cultures to achieve the indicated final concentrations. For each compound and experimental system, the final DMSO concentration was matched to that corresponding to the highest drug concentration tested, and control cultures received the corresponding DMSO concentration in the absence of compound.

In the delayed-treatment paradigm, seeded aggregation was initiated by adding mouse aSyn PFF at DIV13 (D0) in primary neuronal culture. After 7 days (DIV20, D7), compounds were added using the same preparation strategy and maintained for an additional 6 days (DIV26, D13). DMSO controls were matched to the highest DMSO concentration used for each compound in this paradigm. This paradigm was used to assess compound effects on established aSyn pathology rather than on the initiation of aggregation.

To assess PFF internalization, processing, and clearance independently of seeded aggregation, parallel experiments were performed in primary hippocampal neurons derived from aSyn knockout (aSyn KO) mice. In these experiments, aSyn KO neurons were treated at DIV13 with mouse aSyn PFF in the presence of DMSO, 6 µM CI-994, 10 µM AS-252424, or 10 µM CGP-52411. Cells were harvested after 1 (D1), 3 (D3), or 7 (D7) days of treatment for biochemical analysis.

For HCA, primary neurons were cultured and treated in poly-L-lysine-coated black, clear-bottom 96-well plates, and human iDA were cultured on glass coverslips. For biochemical analyses, including Western blotting of detergent-insoluble fractions, neurons were cultured and treated in poly-L-lysine-coated 6-well plates, with two wells analyzed per condition. In all experiments, cultures were maintained without medium exchange following PFF exposure until the experimental endpoint. All experiments were performed using a minimum of three independent biological replicates, each derived from a separate primary neuronal culture preparation. For high-content imaging analyses, three wells per condition were analyzed as technical replicates within each independent biological replicate, as detailed in the corresponding Fig. legends.

### Immunocytochemistry and high-content imaging analysis of aSyn PFF-treated primary neuronal cultures

Following treatment with mouse aSyn PFF and small-molecule modulators, primary hippocampal neurons cultured in black, clear-bottom 96-well plates were washed twice with PBS and fixed with 4% formaldehyde (Sigma-Aldrich, F8775) for 20 min at room temperature. Fixed cells were permeabilized and blocked using our standard immunocytochemical procedures prior to antibody incubation (*76*). Endogenous level of aSyn was detected with SYN-1 antibody (1/1000, BD Biosciences, 610787). Pathological aSyn accumulation was detected using a phosphorylation-specific antibody directed against pS129(1/1000, clone 81A, Biolegend, 825701), which selectively labels aggregated aSyn species. Neuronal morphology and compartmental organization were visualized by co-staining for MAP2 (1/2000, Abcam, 92434), allowing delineation of neuronal cell bodies and neuritic processes. Neuronal nuclei were identified using NeuN (1/5000, Abcam, ab177487) immunoreactivity, and all nuclei present in the mixed neuronal-glial cultures were counterstained with DAPI. Primary antibodies were detected using the following secondary antibodies: donkey anti-chicken Alexa Fluor 488 (1/1000, Jackson ImmunoResearch, 703-545-155), donkey anti-rabbit Alexa Fluor 568 (1/1000, Thermo Fisher Scientific, A-10042), and donkey anti-mouse Alexa Fluor 647 (1/1000, Thermo Fisher Scientific, A-31571).

For aSyn PFF-treated primary mouse hippocampal neurons, images were acquired using an automated wide-field high-content imaging system (INCell Analyzer 2200, GE Healthcare) equipped with a 10× objective. High-content acquisition was performed under identical imaging settings across conditions to enable quantitative comparisons. Each experimental condition was analyzed across three independent biological replicates, each derived from a separate primary neuronal culture preparation. For each biological replicate, three technical replicate wells were imaged per condition, and nine non-overlapping fields of view were collected per well using identical acquisition settings. For aSyn PFF-treated human iDA neurons, imaging was performed using an inverted Zeiss LSM700 confocal microscope. For each independent biological replicate, a minimum of three non-overlapping fields of view were acquired per condition using identical acquisition parameters.

Images were analyzed using a predefined segmentation and analysis pipeline (*76*, *79*). This multiparametric staining strategy enabled quantitative discrimination of pS129-positive aSyn pathology within neuronal somata and MAP2-positive neuritic processes, while simultaneously assessing neuronal survival (NeuN⁺/DAPI⁺), neuritic network integrity (MAP2-positive area), and non-neuronal cell populations (DAPI⁺/NeuN⁻). These parallel measurements allowed aggregation-specific effects to be distinguished from changes in cellular composition or morphology.

### Biochemical analysis of PFF internalization and aSyn pS129 pathology

Following treatment with mouse aSyn PFF and DMSO or compounds, WT or aSyn KO primary hippocampal neurons were harvested for biochemical analyses. Neuronal cultures were lysed in Tris-buffered saline (TBS; 50 mM Tris, 150 mM NaCl, pH 7.5) containing 1% Triton X-100 and supplemented with a protease inhibitor cocktail (Roche, 4693132001), 1 mM phenylmethylsulfonyl fluoride (PMSF, Sigma-Aldrich, P7626), and phosphatase inhibitor cocktails 2 and 3 (Sigma-Aldrich, P5726 and P0044). Cell lysates were subjected to brief probe sonication (10 pulses of 0.5 s at 20% amplitude) using a fine-tip sonicator and incubated on ice for 30 min to ensure efficient solubilization. Lysates were then centrifuged at 100,000 × g for 30 min at 4 °C to separate detergent-soluble and detergent-insoluble material. The supernatant, corresponding to the Triton X-100-soluble fraction, was collected. The pellet was resuspended in fresh 1% Triton X-100/TBS, sonicated as described above, and centrifuged again under the same conditions. Following removal of the supernatant, the resulting pellet, enriched in detergent-insoluble material, was solubilized in 2% SDS/TBS containing protease and phosphatase inhibitors. This insoluble fraction was further homogenized by probe sonication (15 pulses of 0.5 s at 20% amplitude).

Protein concentrations in both soluble and insoluble fractions were determined using the bicinchoninic acid (BCA) assay (Thermo Fisher Scientific, 23222). Samples were subsequently mixed with Laemmli sample buffer (final composition: 4% SDS, 40% glycerol, 0.05% bromophenol blue, 0.252 M Tris-HCl, pH 6.8, and 5% β-mercaptoethanol) and run on 16% Tricine gels, then immunoblotted as previously described (*30*, *76–79*). Western blot analyses were performed to detect total aSyn species (including monomeric ∼15 kDa, truncated ∼12 kDa, and high-molecular-weight assemblies) using SYN-1 antibody (1/1000, BD Biosciences, 610787) and pS129 aSyn (1/10’000, Abcam, MJF-R13, ab168381). Primary antibodies were detected using the following secondary antibodies: goat anti-rabbit Alexa Fluor 800 (1/20’000, LI-COR Biosciences, 926-32211), and goat anti-mouse Alexa Fluor 680 (1/20’000, LI-COR Biosciences, 926-68070). Band intensities were quantified using Image Studio 6.0 software (LI-COR Biosciences) and normalized to actin levels. Densitometric values are expressed as fold change relative to the control condition, as specified in the corresponding figure legends. Each data point represents an independent biological replicate from primary neuronal cultures prepared on different days. Due to gel capacity constraints, samples from independent experiments were processed on separate gels and membranes rather than run simultaneously. This experimental design is reflected in the presentation of the quantitative analyses.

### Statistical analysis of aSyn-related experiments

Statistical analyses were performed using GraphPad Prism software (RRID: SCR_002798). All quantitative data were derived from at least three independent biological experiments. Group comparisons were carried out using one-way analysis of variance (ANOVA), followed by Tukey’s post hoc test to correct for multiple comparisons, as specified in the corresponding Fig. legends. Statistical significance was defined as P < 0.05.

## Supplementary Information Inventory

### Supplementary Figures

There are 7 Supplementary figures. Their relationship to the main figures are indicated in the figure legends.

### Supplementary Tables

There are 2 Supplementary tables providing Quality assessment of plates used in the drugs screening and the list of compounds included in the secondary screen and their assigned mechanisms of action (MOA).

### Supplementary Files

There are two Supplementary files (not included in this document) providing the list of compounds of the in-house kinase inhibitors and drug-repurposing libraries.

### Supplementary file S1. (separate file)

List of the compounds in the in-house kinase inhibitors library.

### Supplementary file S2. (separate file)

List of the compounds in the in-house drug repurposing library.

---

**Fig. S1.**
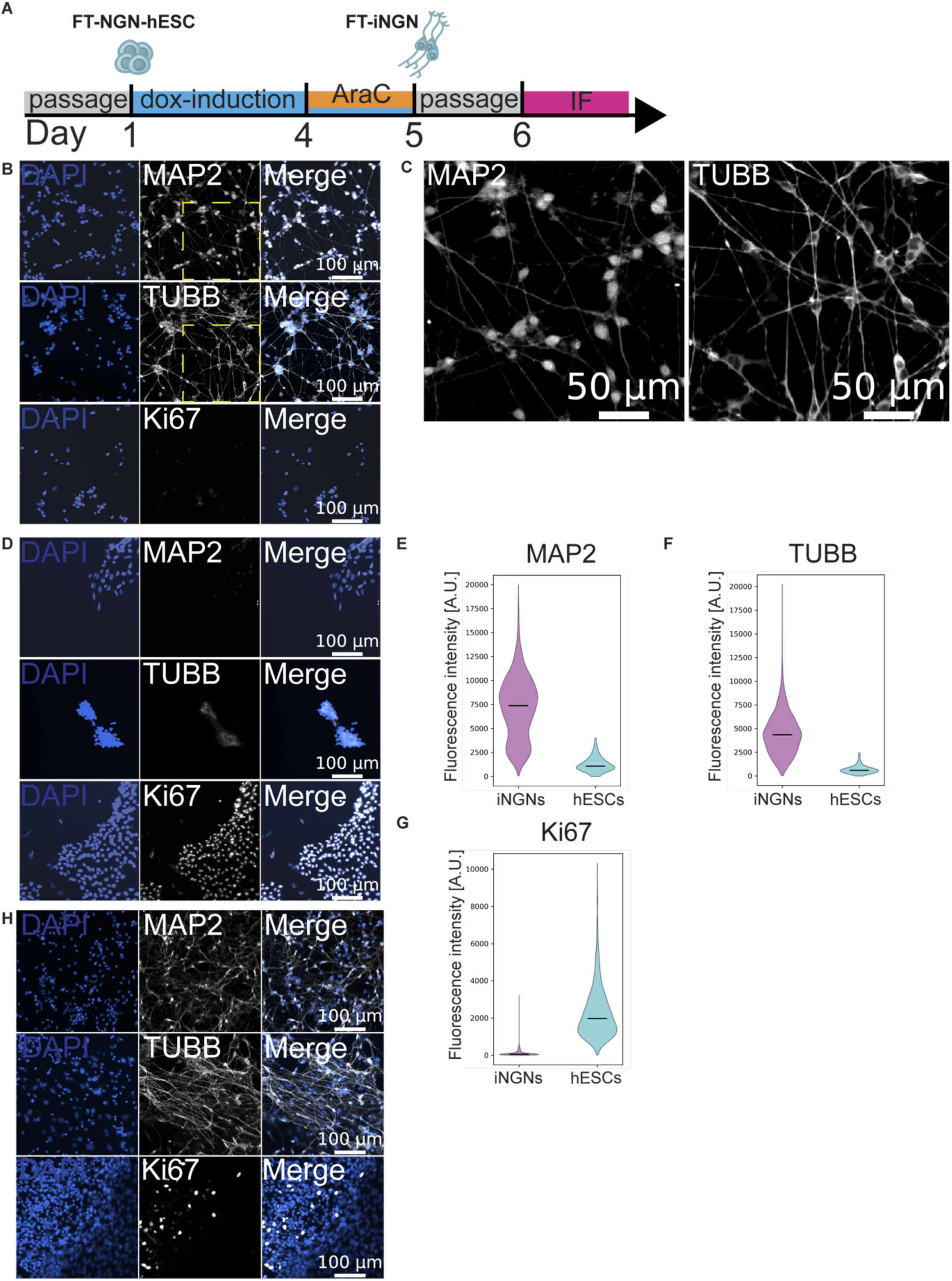
related to Fig. 1. Validation of the neuronal differentiation models. **A.** Experimental workflow of differentiating human embryonic stem cells (FT-NGN-hESCs) to dox-induced neurons (FT-iNGNs). **B-D**: Representative immunofluorescence labeling images of the neuronal markers TUBB, MAP2 and the proliferation marker Ki67 in FT-iNGNs (**B**) made as described in (**A**) (higher magnifications shown in **C**) or in undifferentiated FT-NGN-hESCs (**D**). Nuclei were stained with DAPI (blue). **E-G.** Quantification of immunofluorescence labeling intensity from (**B**) and (**D**) of FT-NGN-hESCs (cyan) and FT-iNGNs (magenta) for MAP2 (**E**), TUBB (**F**), Ki67 (**G**). Violin plot represents per-cell data from two replicates. N> 100 cells. Black line: median fluorescence intensity. **H.** Representative immunofluorescence labeling images of the neuronal markers TUBB, MAP2 and the proliferation marker Ki67 in neuron-enriched culture based on 2D differentiation protocol and derived from FT-hESCs. Nuclei were stained with DAPI (blue).

**Fig. S2.**
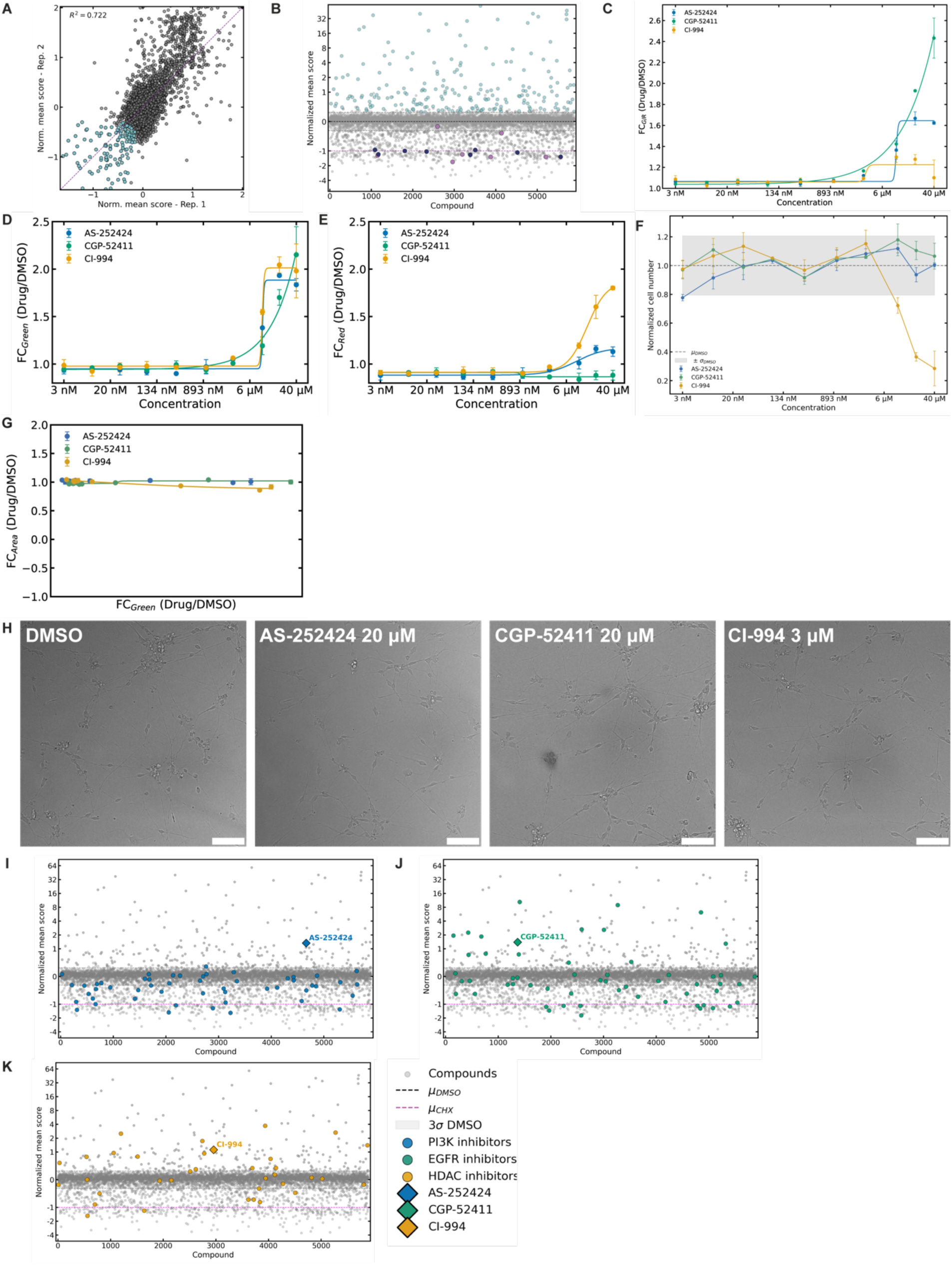
related to Fig. 2. Primary and secondary screening reveal protein turnover modulators. **A.** Linear correlation between normalized scores of 5481 tested compounds (gray), including hits (cyan), in two replicates from the primary screen. At least 800 cells per compound were measured. Values for 1^st^–99^th^ percentiles are visualized, omitting compounds that didn’t pass the z’-factor quality control. **B.** Normalized inverted mean values of drugs tested (N=2) in the primary screen with annotated known synthesis and proteasome inhibitors, in orchid and navy, respectively. Compounds (gray) were annotated as hits (cyan) when they scored above 3 standard deviations (s) of the DMSO control. The inverted values are displayed, where the value for CHX is-1 (dashed magenta line). At least 800 cells per compound per replicate were measured. Synthesis Inhibitors: Harringtonine, Sal003, Puromycin (Dihydrochloride), NH125, Emetine (dihydrochloride hydrate), Salubrinal, Homoharringtonine. Proteasome inhibitors: MG-132, MLN2238, Oprozomib, Carfilzomib, delanzomib, ONX-0914, MLN9708, ixazomib-citrate. **C-E.** Dose-response curve of the selected compounds in the secondary screen for the G/R ratio (**C**) and Green (**D**) or Red fluorescence (**E**). Fold change (FC) of a compound to the mean of the vehicle (DMSO). The sigmoidal curve (solid line) was fitted to the mean of the data point from N=2, error bar: standard deviation. **F.** Cytotoxicity of the selected compounds in the secondary screen. Cell numbers were normalized to mean cell numbers in the vehicle (DMSO) condition. Each dot represents a replicate (N=2, at least 800 cells per replicate were measured), error bar: standard deviation, dashed line: vehicle mean value, gray-colored window: 3 standard deviation values in each direction. **G.** FC of nucleus area (calculated based on the segmentation mask) to FC of normalized green fluorescence, for the selected compounds in the secondary screen, normalized to the mean vehicle (DMSO) compared. (N=2, at least 800 cells per replicate were measured), error bar: standard deviation. **H**. Representative brightfield images of FT-iNGNs treated for 24 h with DMSO or 20 µM AS-252424, 20 µM CGP-52411 or 3 µM CI-994. Scale bar: 100 µm. **I-K.** Primary screen data, as explained in (**B**), highlighting drugs from the same class as the selected compounds (diamond) are shown in (**I**) AS-252424 and PI3K inhibitors (blue); (**J**) CGP-52411 and EGFR inhibitors (green); (**K**) CI-994 and HDAC inhibitors (yellow).

**Fig. S3.**
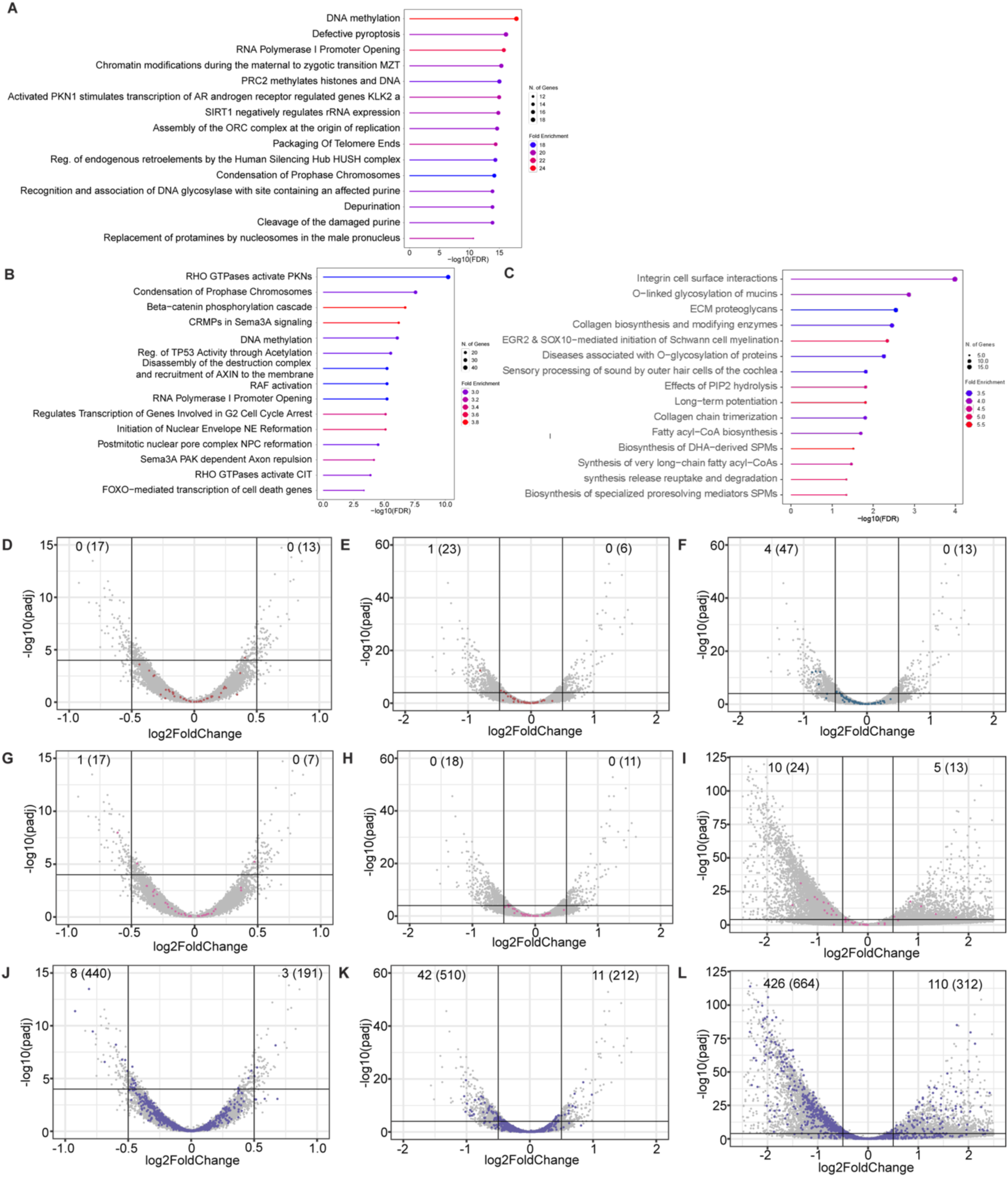
related to Fig. 3. Lack of a distinct signature among downregulated genes and EGFR/PI3K/mTOR pathway alterations. Reactome Knowledgebase enrichment analyses of significantly enriched pathways downregulated upon CGP-52411 treatment (**A**), downregulated (**B**) and upregulated (**C**) upon CI-994 treatment. Expression changes of PI3K pathway-associated genes upon 24 h treatment with AS-252424 (**D**) and CGP-52411 (**E**). Expression changes of EGFR pathway-associated genes upon 24 h treatment with CGP-52411 (**F**). Expression changes of mTOR-associated genes upon 24 h treatment with AS-252424 (**G**), CGP-52411 (**H**) and CI-994 (**I**). (**J-L**) Expression changes of neurogenesis-related genes upon 24h treatment with AS-252424 (J), CGP-52411 (K) and (L) CI-994. Number of significantly downregulated or upregulated genes (log2FC>|0.5|, padj<0.0001) as well as genes with log2FC>|0.1| (number between brackets) are displayed on the upper corners of each plots. Data are the results of two independent replicates.

**Fig. S4.**
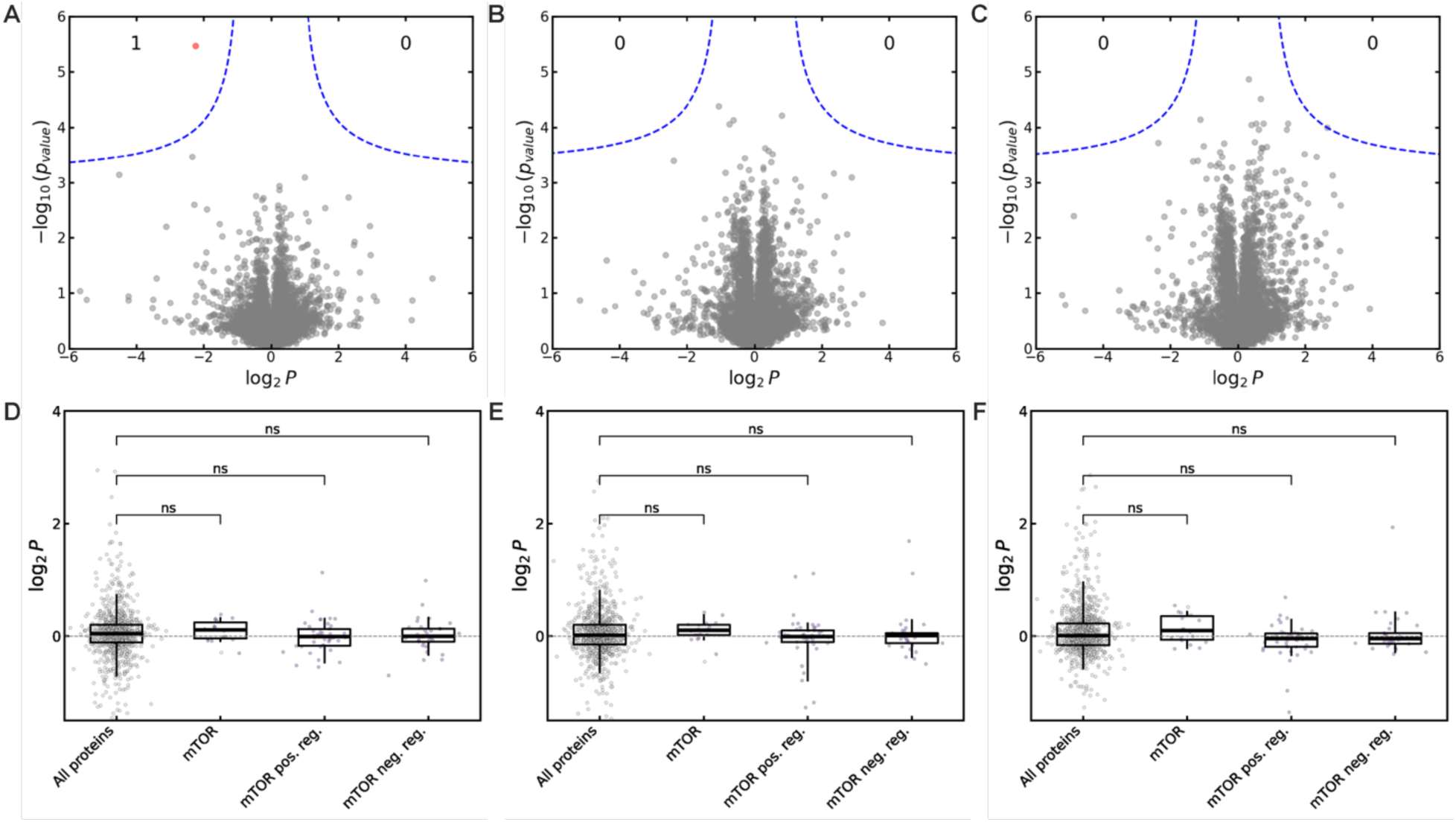
related to Fig. 4. AS-252424, CGP-52411 and CI-994 do not affect levels of all proteins or mTOR signaling-related proteins. Label-free proteome quantification for iNGNs upon 24 h treatment with AS-252424 (**A**), CGP-52411 (**B**) and CI-994 (**C**), for 5713 proteins common to control and treated conditions. **D-F**. Comparison of differential expression for mTOR signaling and positive/negative regulator (pos. reg./neg. reg.) proteins against all other proteins for AS-252424 (**D**), CGP-52411 (**E**) and CI-994 (**F**). Each dot represents a mean of N=3; boxes: interquartile range; horizontal line: median; dashed line: median of all proteins; vertical lines: 5^th^-95^th^ percentiles; ns - non-significant.

**Fig. S5.**
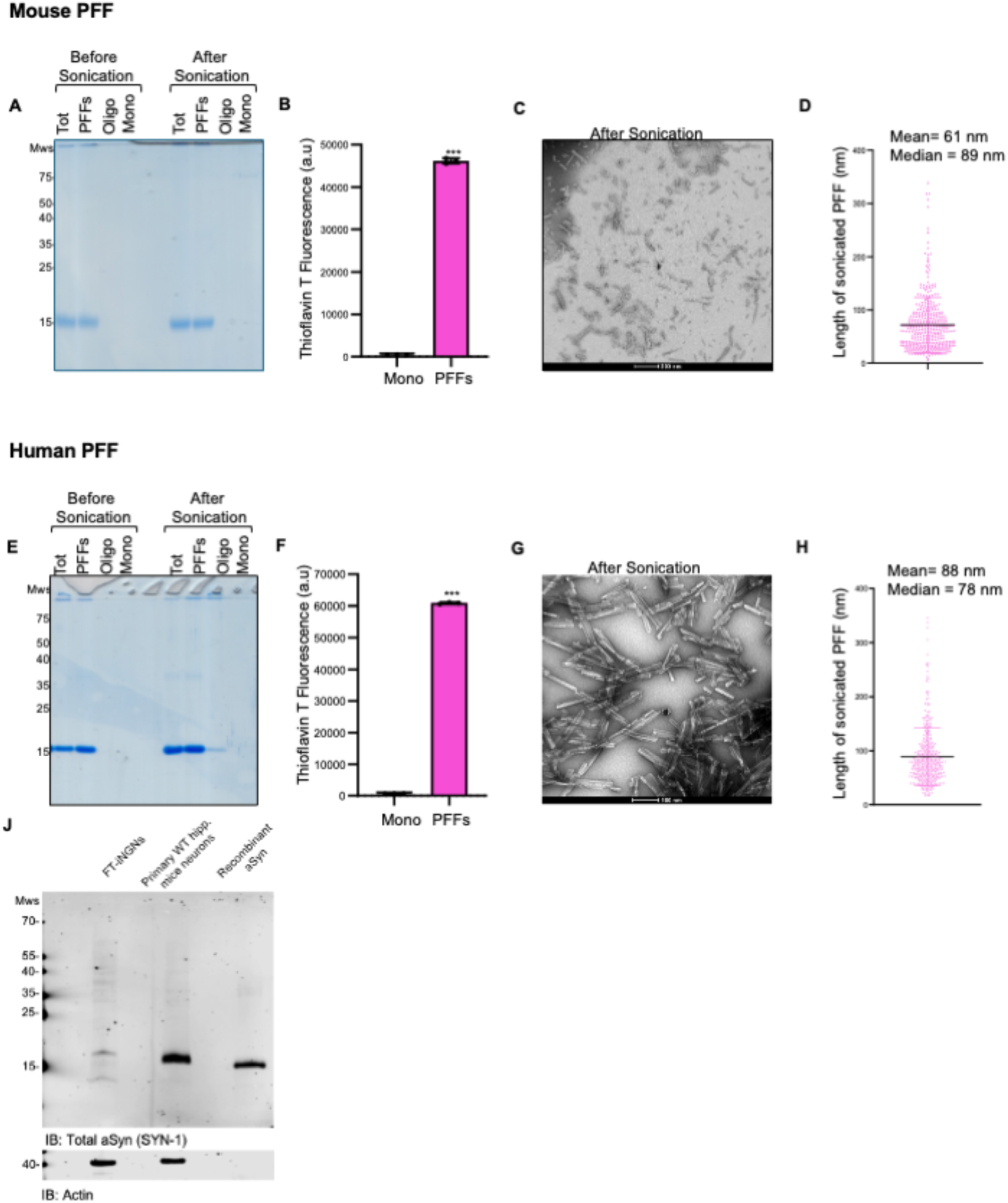
related to Fig. 5. Mouse and human aSyn PFF consist of β-sheet-rich amyloid fibrils and short seeds suitable for neuronal seeding assays. Mouse (m) and human (h) recombinant aSyn fibrils were generated and characterized prior to use in neuronal seeding experiments. **A, E.** Coomassie-stained SDS-PAGE showing mouse (**A**) or human (**E**) aSyn monomer and PFF before and after sonication. Monomeric aSyn migrates at ∼15 kDa, whereas fibrillar preparations display characteristic higher molecular weight species that are retained at the top of the gel (stacking region), consistent with large insoluble aggregated aSyn assemblies. **B, F.** Thioflavin T (ThT) fluorescence assay demonstrating robust β-sheet-rich amyloid formation in mouse (**B**) and human (**F**) PFF preparations compared to monomeric aSyn. Data represent mean ± s.d. from n = 3 technical replicates. Statistical significance was assessed by an unpaired two-tailed Student’s t-test. **C, G.** Representative transmission electron microscopy (TEM) image of sonicated mouse (**C**) or human (**G**) aSyn PFF, revealing short fibrillar species. **D, H.** Quantification of fibril length distribution for mouse (**D**) and human (**H**) aSyn PFF after sonication. **D, H.** Each dot represents an individual fibril measured from TEM micrographs. Pink line: mean; black line: median. **J.** Western blotting against aSYN in FT-iNgns, primary mouse neurons, and on recombinant aSyn.

**Fig. S6.**
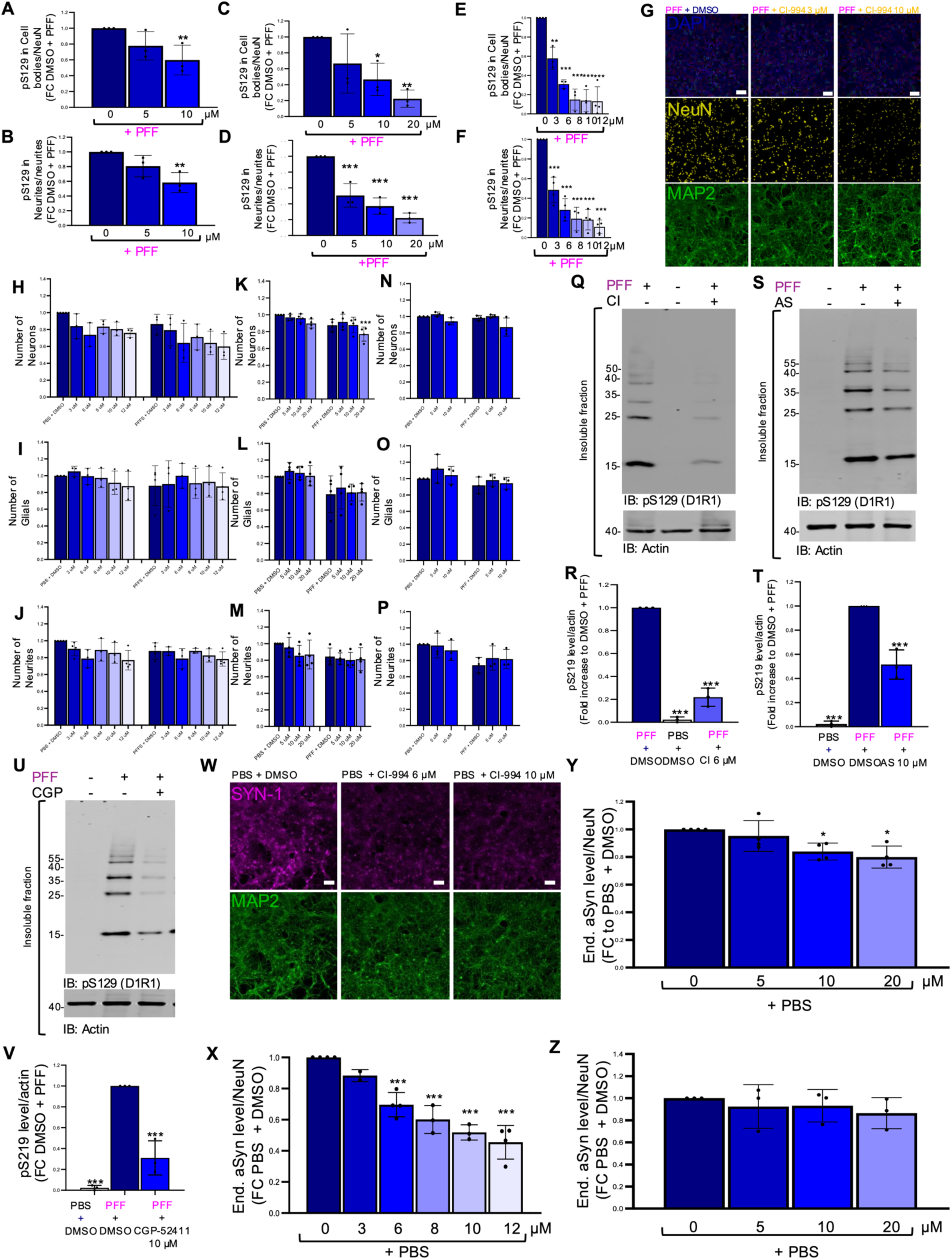
related to Fig. 5. CI-994, AS-252424, and CGP-52411 suppress seeded aSyn pathology during co-treatment with mouse aSyn PFF in primary mouse neurons without inducing cytotoxicity. **A-F**. Quantification of the impact of CGP-52411 (**A-B**), AS-252424 (**C-D**) and CI-994 (**E-F**) on pS129-positive aSyn pathology during co-treatment by HCA in the soma (**A, C** and **E**) or in neurites, quantified within MAP2-positive processes (**B, D** and **F**). **G.** Representative immunocytochemistry images of primary hippocampal neurons exposed to aSyn PFF and co-treated with DMSO or CI-994 (3-10 µM). Neuronal nuclei were labeled with NeuN (yellow), neuronal somata and neuritic processes with MAP2 (green), and all nuclei (neuronal and non-neuronal) with DAPI (blue). Scale bar: 50 µm. **H-P.** Effect of drugs on cell survival. Primary mouse hippocampal neurons were treated with PBS or aSyn PFF in the presence of DMSO or increasing concentrations of CI-994 (**H-J**), AS-252424 (**K-M**), or CGP-52411 (**N-P**), and cellular parameters were quantified by high-content analysis at D10 post-treatment. **H, K, N.** Neuronal counts quantified as NeuN⁺/DAPI⁺ objects. **I, L, O.** Glial cell counts quantified as DAPI⁺/NeuN⁻ objects. **J, M, P.** Neuritic network density quantified from MAP2-positive processes. **Q, S, U.** WB analysis of detergent-insoluble fractions from neurons treated with PBS or PFF in the presence of DMSO or (**Q**) CI-994 (6 µM), (**S**) AS-252424 (10 µM), (**U**) CGP-52411 (10 µM), immunoblotted for pS129-aSyn (D1R1). **R, T, V.** Quantification of insoluble pS129-aSyn levels shown in (**Q,S,U**) for (**R**) CI-994 (6 µM), (**T**) AS-252424 (10 µM), (**V**) CGP-52411 (10 µM), expressed as fold change relative to PFF + DMSO controls, normalized to actin. **W.** Representative immunocytochemistry images showing total endogenous aSyn levels detected using the anti-SYN-1 antibody (magenta) in PBS-treated neurons co-treated with DMSO or CI-994 (6 or 10 µM). An anti-MAP2 antibody (green) labels neuronal processes. **X-Z.** Concentration-dependent changes in endogenous aSyn levels upon CI-994 (**X**), AS-252424 (**Y**) and CGP-52411 treatment (**Z**). All quantitative data (**C-F, H-P, R, T, V, X-Z**): mean ± s.d. from a minimum of N = 3 biological replicates and n = 3 technical replicates, each corresponding to an independent primary neuronal culture preparation. *P < 0.05; **P < 0.01; ***P < 0.001.

**Fig. S7.**
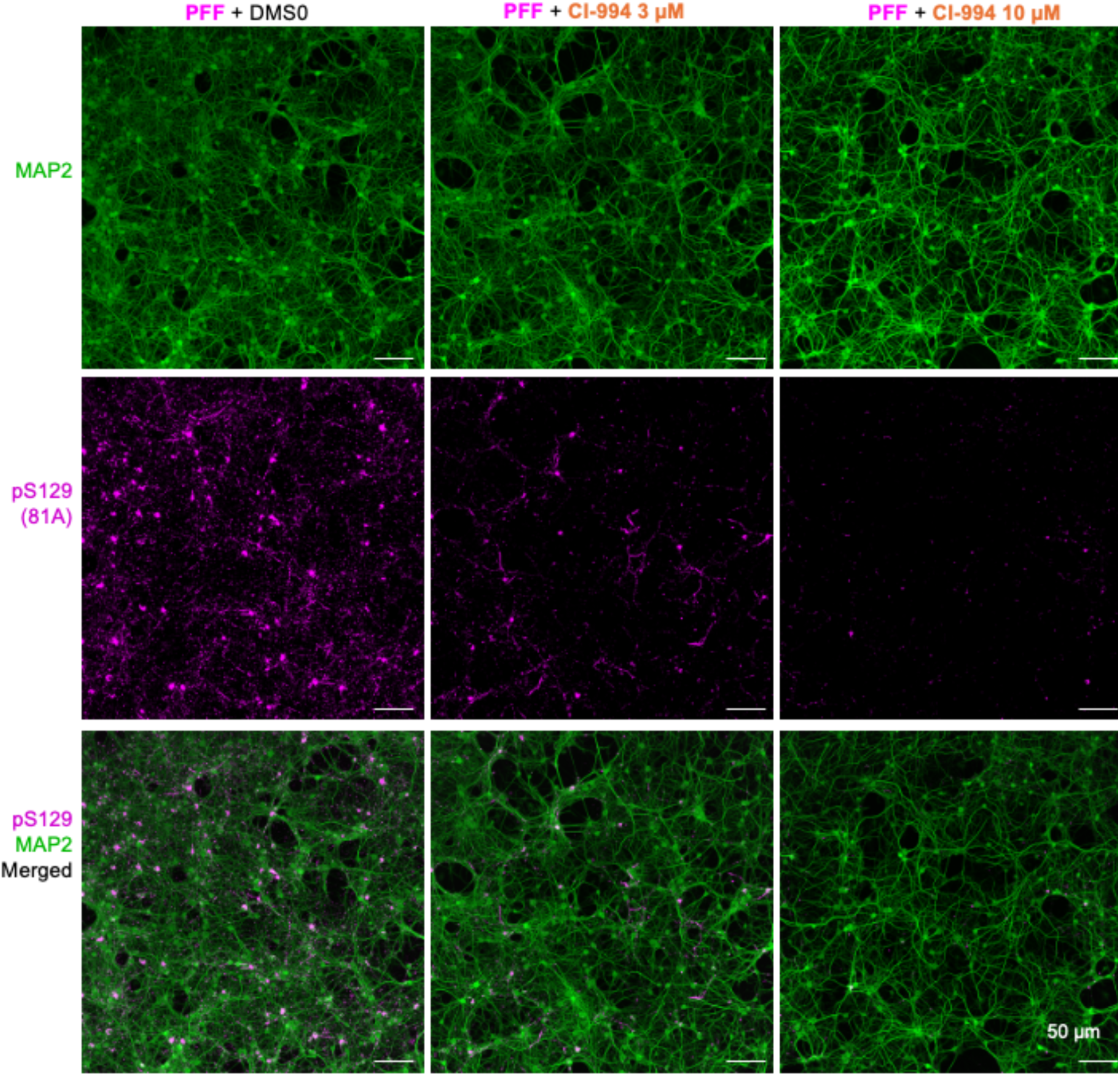
related to Fig. 5B. Enlarged insets of the representative immunocytochemistry images of neurons treated with PFFs and co-treated with DMSO or CI-994 (3-10 µM). Pathological aSyn aggregates were detected using a phosphorylation-specific pS129 antibody (81A; magenta), and neuronal soma and neurites were detected with MAP2 (green). Merged images are shown below.

**Fig. S8.**
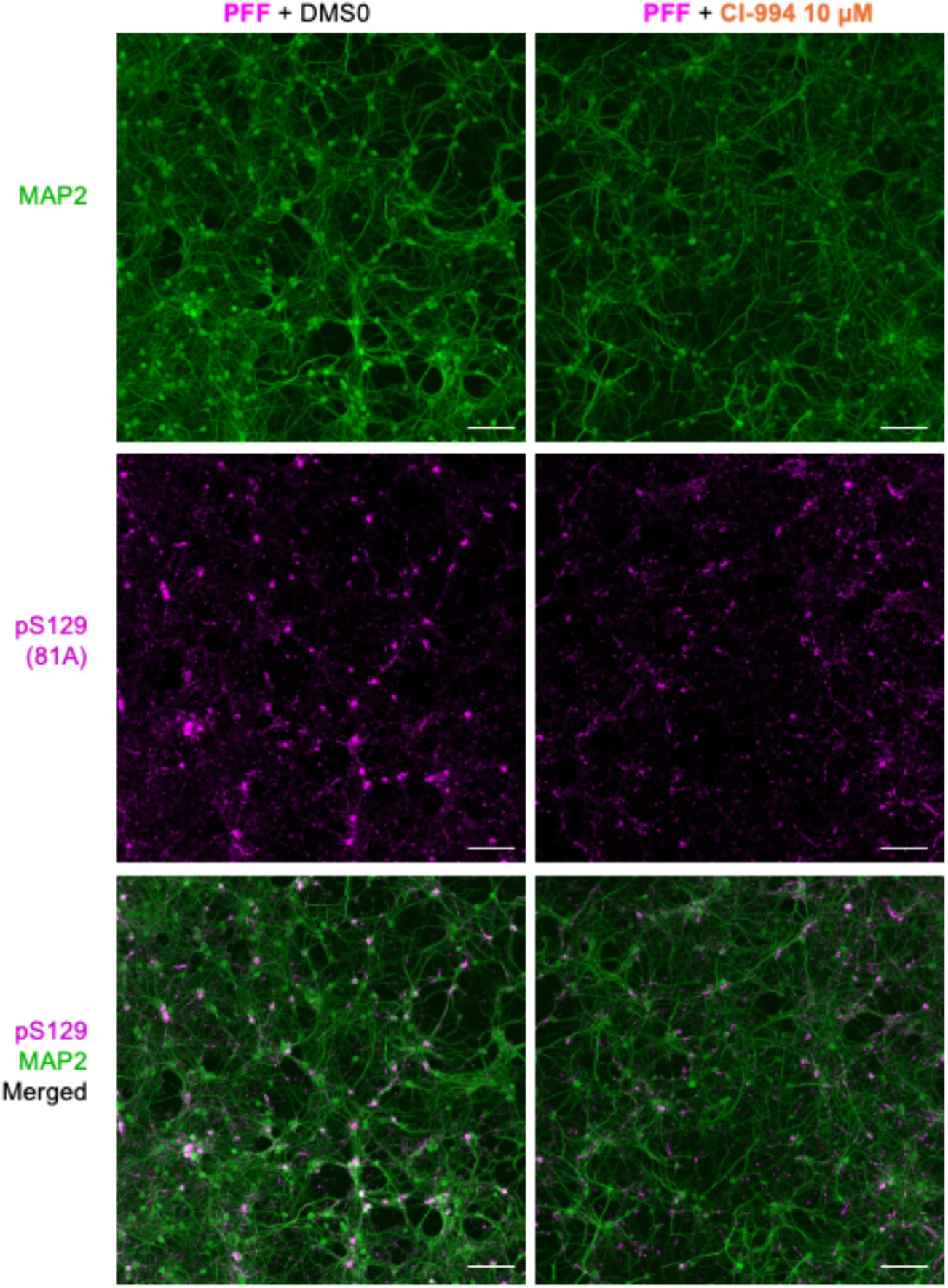
related to Fig. 5G. Enlarged insets of the representative images of PFF-treated neurons after delayed treatment with DMSO or CI-994 (10 µM). Pathological pS129-positive aggregates were detected with 81A (magenta), and neurons with MAP2 (green). Merged images are shown in the bottom panel. Scale bar: 50 µm.

**Fig. S9.**
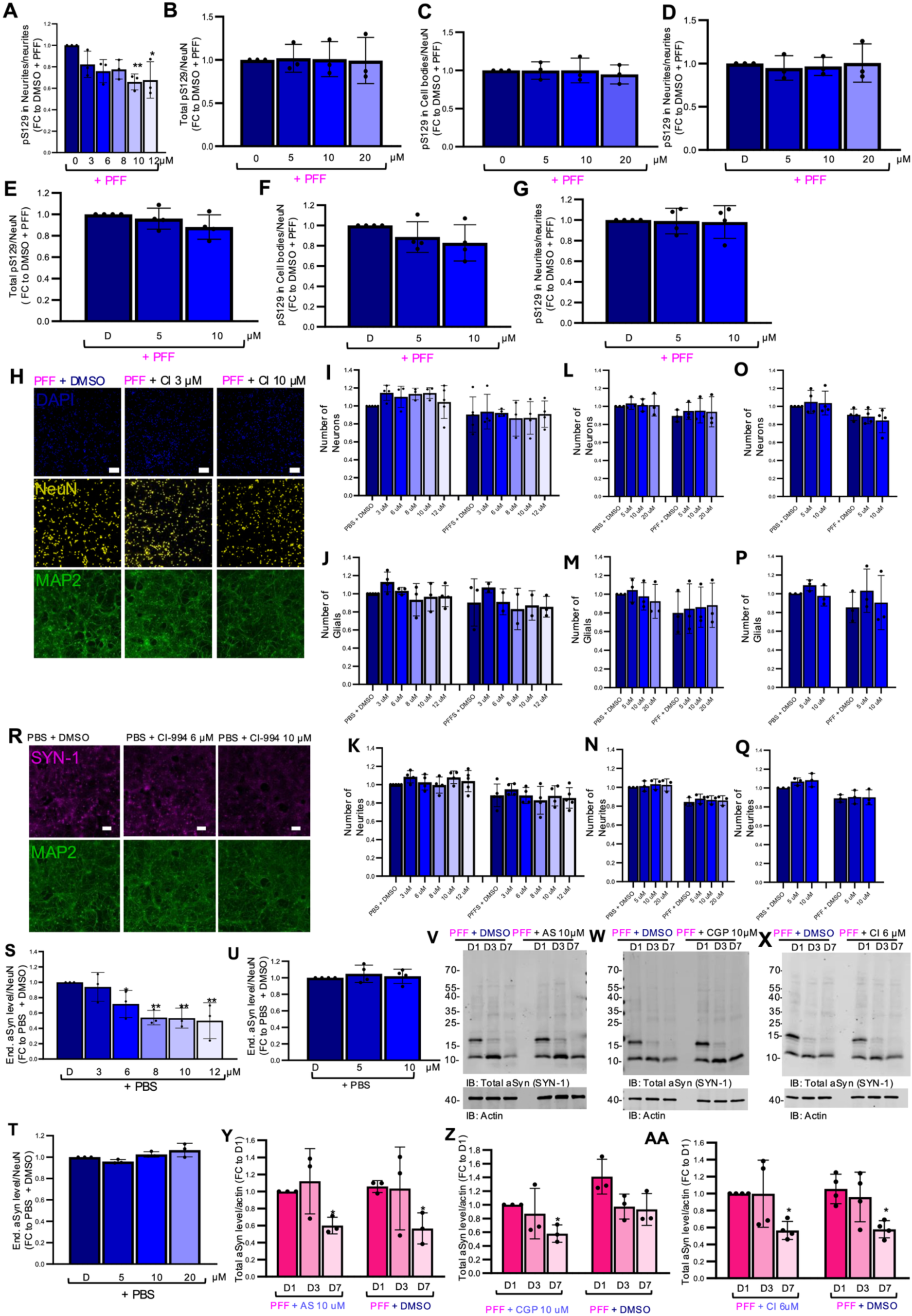
related to Fig. 5. CI-994 acts on pre-established aSyn pathology in primary mouse neurons. CI-994, AS-252424 and CGP-52411 do not affect fibril uptake. **A-G.** Quantification of pS129 pathology by high-content analysis under delayed-treatment conditions, shown and expressed as fold change relative to PFF + DMSO controls. **A.** Neuritic pS129 pathology quantified within MAP2-positive processes upon CI-994 treatment. Total pS129 signal (**B**), somatic pS129 pathology within NeuN-positive neuronal cell bodies (**C**), and neuritic pS129 pathology within MAP2-positive processes (**D**) upon AS-252424 treatment. Total pS129 signal (**E**), somatic pS129 pathology within NeuN-positive neuronal cell bodies (**F**), and neuritic pS129 pathology within MAP2-positive processes (**G**) upon CGP-52411 treatment. **H.** Representative immunocytochemistry images of PFF-treated primary hippocampal neurons following delayed treatment with DMSO or CI-994 (3-10 µM). Neuronal nuclei were labeled with NeuN (yellow), neuronal somata and neuritic processes with an anti-MAP2 antibody (green), and nuclei with DAPI (blue). Scale bar: 50 µm. **I-Q.** Primary mouse hippocampal neurons were treated with PBS or aSyn PFF in the presence of DMSO or increasing concentrations of CI-994 (**I-K**), AS-252424 (**L-N**), or CGP-52411 (**O-Q**), and cellular parameters were quantified by high-content analysis at D10 post-treatment. All values are expressed as fold change relative to the corresponding PBS + DMSO condition. **I, L, O.** Neuronal counts quantified as NeuN⁺/DAPI⁺ objects. **J,M,P.** Glial cell counts were quantified as DAPI⁺/NeuN⁻ objects. **K,M,Q.** Neuritic network density quantified from MAP2-positive processes. **R.** Representative immunocytochemistry images showing endogenous aSyn levels detected using an anti-SYN-1 antibody (magenta) in naïve PBS-treated neurons exposed to DMSO or CI-994 (6 or 10 µM) under delayed-treatment conditions. MAP2 (green) labels neuronal processes. Scale bar: 50 µm. **S-U.** Quantification of endogenous aSyn levels measured in PBS-treated neurons under delayed-treatment conditions for CI-994 (**S**), AS-252424 (**T**), and CGP-52411 (**U**), expressed as fold change relative to PBS + DMSO controls. **V-X.** WB analysis of internalized PFF in aSyn KO primary neurons exposed to PFF in the presence of DMSO or (**V**) AS-252424 (10 µM), (**W**) CGP-52411 (10 µM), (**X**) CI-994 (6 µM) for 1, 3, or 7 days. Blots were probed for total aSyn (SYN-1). Actin: loading control. **Y-AA.** Quantification of total aSyn signal shown in (**V-X**) for (**Y**) AS-252424 (10 µM), (**Z**) CGP-52411 (10 µM), (**AA**) CI-994 (6 µM), expressed as fold change relative to D1 DMSO controls. All quantitative data (**A-G**, **I-Q**, **S-U**, **Y-AA**): mean ± s.d. from a minimum of N = 3 biological replicates and for images also n = 3 technical replicates, each corresponding to an independent primary neuronal culture preparation. *P < 0.05; **P < 0.01; ***P < 0.001.

**Fig. S10.**
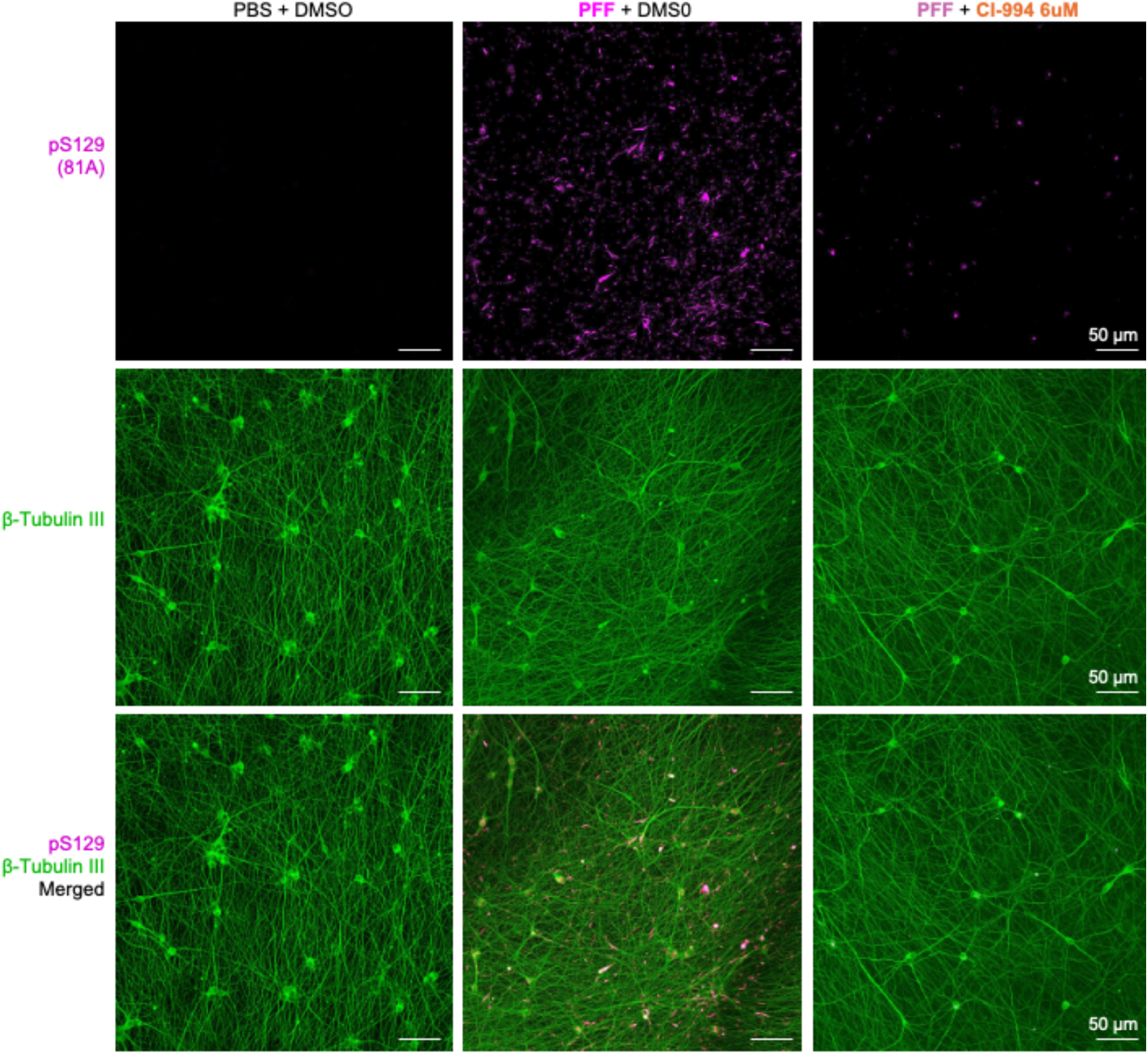
related to Fig. 5K. Enlarged insets of the representative images of iDA neurons treated with PBS + DMSO, PFF + DMSO or PFF + 6 µM CI-994. Pathological aggregates were detected using pS129 (81A; magenta), and neuronal processes were detected using βIII-tubulin (green). Merged images are shown in the bottom panel.

**Table S1.** Quality assessment of plates used in the drugs screening. z^’^-factor per plate was calculated based on vehicle DMSO and inverse CHX controls present in two columns each in each plate. *: rejected plate due to the negative value of z’-factor.

| Library | PlateID | z'-factor |
| --- | --- | --- |
| Kinase Inhibitors | BSF026213 | 0.53516626 |
|  | BSF026214 | 0.44394987 |
| Prestwick Chemical Library | BSF026206 | 0.45507371 |
|  | BSF026330 | 0.74472376 |
|  | BSF026208 | 0.56841701 |
|  | BSF026331 | 0.69577254 |
|  | BSF026210 | 0.60733983 |
|  | BSF026332 | 0.67190848 |
|  | BSF026212 | 0.64060823 |
|  | BSF026333 | 0.72789954 |
| Drug Repurposing | BSF026334 | 0.69280599 |
|  | BSF026335 | 0.82527486 |
|  | BSF026336 | -5.1086577* |
|  | BSF026337 | 0.82686056 |
|  | BSF026338 | 0.40906849 |
|  | BSF026339 | 0.83624159 |
|  | BSF026340 | 0.42955448 |
|  | BSF026341 | 0.8450325 |
|  | BSF026342 | 0.43704446 |
|  | BSF026343 | 0.84460223 |
|  | BSF026344 | 0.44873455 |
|  | BSF026345 | 0.76757829 |
|  | BSF026346 | 0.25124765 |
|  | BSF026347 | 0.75615357 |
|  | BSF026348 | 0.44381102 |
|  | BSF026349 | 0.71549242 |
|  | BSF026350 | 0.34023744 |
|  | BSF026351 | 0.74889001 |
|  | BSF026352 | 0.66258991 |
|  | BSF026353 | 0.72961973 |
|  | BSF026354 | 0.71837616 |
|  | BSF026355 | 0.68072345 |
|  | BSF026356 | 0.68194285 |
|  | BSF026357 | 0.7362439 |
|  | BSF026358 | 0.54517743 |
|  | BSF026359 | 0.75307542 |
|  | BSF026360 | 0.69508058 |
|  | BSF026361 | 0.68253974 |
| Secondary screen | BSF026781 | 0.647243365 |
|  | BSF026783 | 0.753063525 |
|  | BSF026785 | 0.54558582 |

**Table S2.**
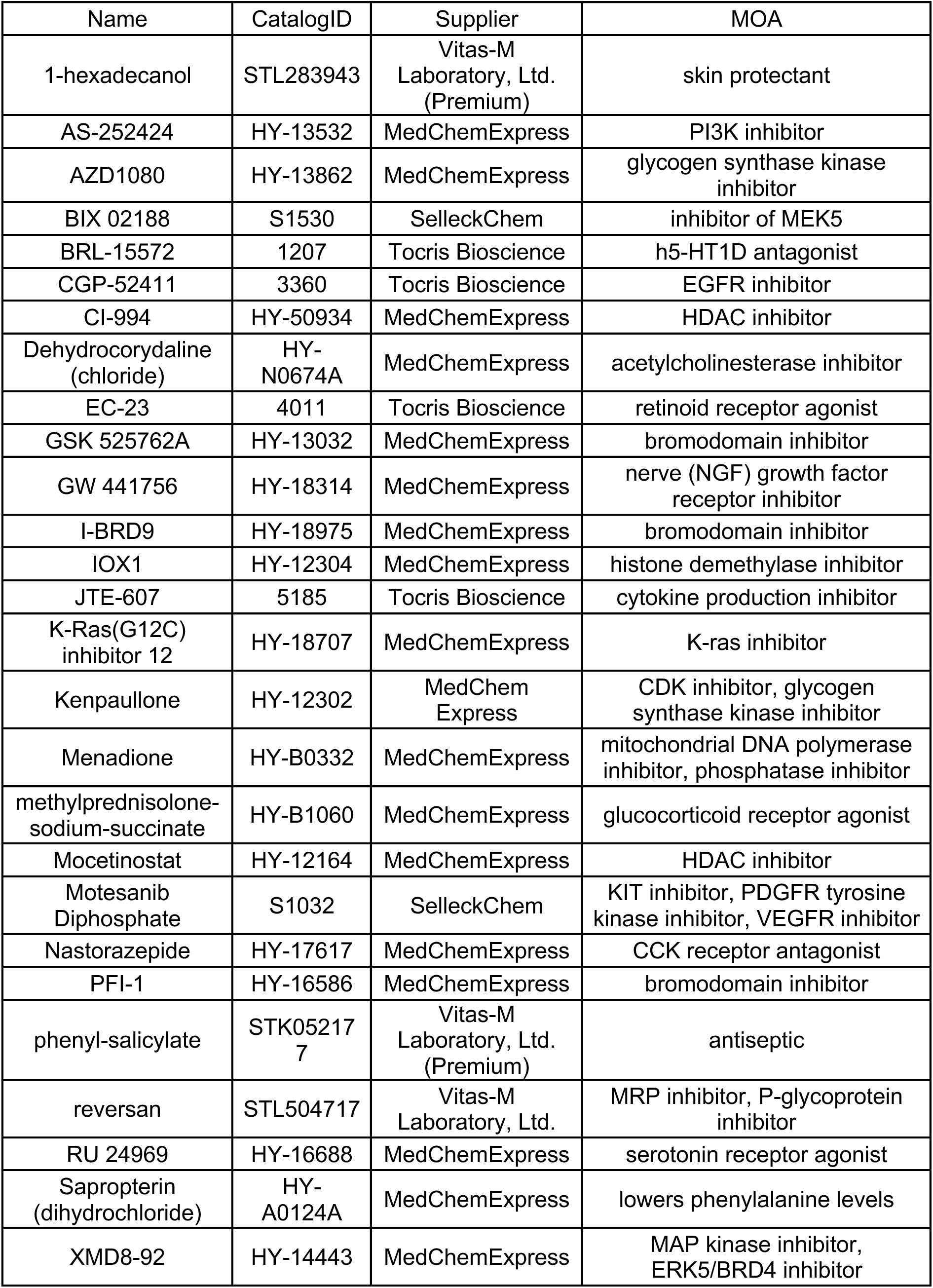
List of compounds included in the secondary screen and their assigned mechanisms of action (MOA). Excluding compounds removed due to precipitation or imaging artifacts.

